# Selectivity filter mutations shift ion permeation mechanism in potassium channels

**DOI:** 10.1101/2023.04.17.537168

**Authors:** Andrei Mironenko, Bert L. de Groot, Wojciech Kopec

## Abstract

Potassium (K^+^) channels combine high conductance with high ion selectivity. To explain this efficiency, two molecular mechanisms have been proposed. The ‘direct knock-on’ mechanism is defined by water-free K^+^ permeation and formation of direct ion-ion contacts in the highly conserved selectivity filter (SF). The ‘soft knock-on’ mechanism involves co-permeation of water and separation of K^+^ by water molecules. With the aim to distinguish between these mechanisms, crystal structures of the KcsA channel with mutations in two SF residues - G77 and T75 - were published, where the arrangements of K^+^ ions and water display canonical soft knock-on configurations. These data were interpreted as evidence of the soft knock-on mechanism in wild-type channels (C. Tilegenova, *et al.*, Structure, function, and ion-binding properties of a K+ channel stabilized in the 2,4-ion–bound configuration. *Proceedings of the National Academy of Sciences* **116**, 16829–16834 (2019)). Here, we test this interpretation using molecular dynamics simulations of KcsA and its mutants. We show that, while a strictly water-free direct knock-on permeation is observed in the wild-type, conformational changes induced by these mutations lead to distinct ion permeation mechanisms, characterized by co-permeation of K^+^ and water. These mechanisms are characterized by reduced conductance and impaired potassium selectivity, supporting the importance of full dehydration of potassium ions for the hallmark high conductance and selectivity of K^+^ channels. In general, we present a case where mutations introduced at the critical points of the permeation pathway in an ion channel drastically change its permeation mechanism in a non-intuitive manner.

**Significance statement:** Potassium (K^+^) channels conduct K^+^ with high permeation rates and ion selectivity. An ongoing debate in the field has been focused on the molecular mechanisms underlying this remarkable efficiency. Here, we performed molecular dynamics simulations of two selectivity filter mutants of a model K^+^ channel to investigate this question. These mutations led to a substantial decrease in conductance and ion selectivity, while accompanied by a shift from water-free K^+^ permeation to co-permeation of water and K^+^. Our findings not only provide a fundamental example of how single point mutations in the selectivity filter can alter the ion permeation mechanism, but also reinforce the notion that water exclusion underlies the remarkable efficiency of K^+^ channels.

## Introduction

Potassium channels (K^+^ channels) enable the permeation of K^+^ ions along their electrochemical gradient across plasma and organelle membranes in almost all organisms. They play an essential role in establishing the membrane potential in living cells, terminating action potentials in excitable cells, as well as in many other physiological processes (1). K^+^ channels are able to conduct K^+^ with very high permeation rates, with conductances reaching hundreds of pS (2). At the same time, they possess exceptional selectivity for K^+^ over other monovalent ions, with K^+^/ Na^+^ permeability ratios reaching ∼100-1000 (3). In the last decades a great effort has been put into determining the ion permeation mechanism that can explain such high K^+^ permeation efficiency, combined with strict K+ selectivity (4).

K^+^ channels possess a highly conserved selectivity filter (SF), usually with a signature sequence TVGYG (5, 6). The SF is the narrowest part of a K^+^ channel pore, and serves as the functional core of the channel. It consists of 4 K^+^ binding sites - S1 to S4 respectively-lined by backbone carbonyls (S1-S4) and threonine hydroxyls (S4 only), pointing toward the permeation pathway (Fig. 1*A*). The K^+^ permeation mechanism in K^+^ channels is defined by specific arrangements of K^+^ ions and, possibly, water molecules inside SF, occuring during ion translocation through the channel. Currently, there are two primary models of K^+^ permeation in K^+^ channels: the ‘soft knock-on’ and ‘direct knock-on’ mechanisms (4). Soft knock-on was proposed following the first structural data: it states that the SF is occupied by 2 K^+^ simultaneously - either in S1 and S3, or S2 and S4 - and by water molecules in the remaining 2 sites (7, 8). This separation of two K^+^ ions by a water molecule was included to prevent seemingly too strong electrostatic repulsion between K^+^ ions, if they would occupy neighboring sites. Accordingly, in the course of a permeation event, given an initial configuration ‘KWKW’ (K stands for a K^+^ ion, and W for a water molecule, in sites S1 to S4 respectively), an ion approaching S4 from the intracellular side ‘knocks on’ the contents of the SF to a configuration ‘WKWK’ (Fig. 1*A*), and vice versa - thus, for each K^+^ ion there is one water molecule permeating the channel. A very different process takes place in the direct knock-on mechanism, as water does not permeate the channel. Rather, K^+^ ions are completely dehydrated in the SF, and direct ion-ion contacts are formed (Fig. 1*A*) (9). This complete dehydration of K^+^ ions in the SF makes ion permeation inherently selective towards K^+^ compared to Na^+^ due to the lower dehydration free energy of the former (10). In addition, the strong electrostatic repulsion arising from the direct ion-ion contacts was proposed to drive fast permeation in K^+^ channels (9). Importantly, both soft and direct knock-on mechanisms follow a strict single-file ion (and water) permeation and an unperturbed SF conformation is assumed during ion permeation.

**Fig. 1.**
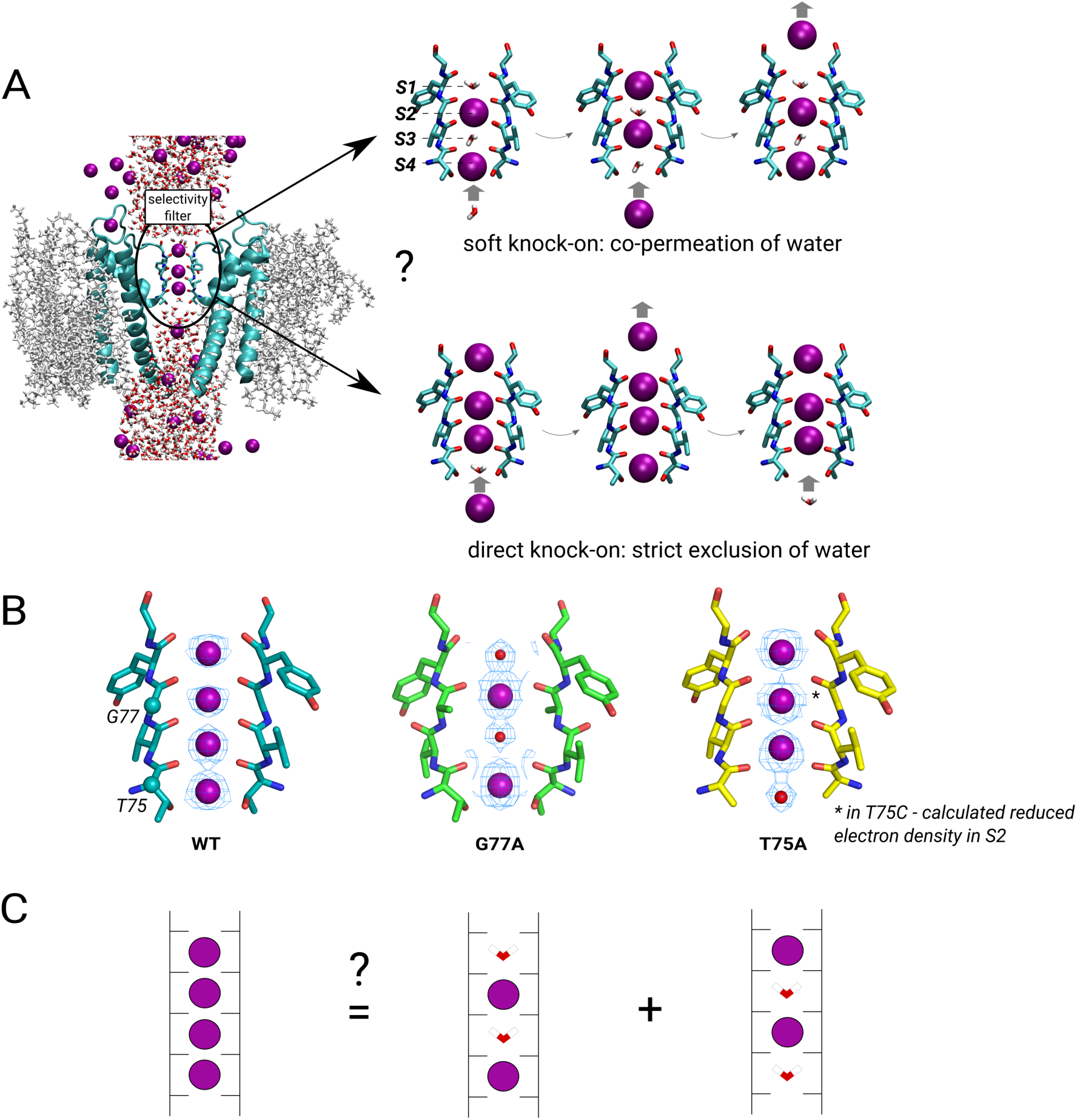
Overview of ion permeation mechanisms in potassium channels. (*A*) Side view of a structure of KcsA embedded in a lipid membrane, with the highly conserved selectivity filter (SF) highlighted. K^+^ ions are shown as purple spheres. On the right, main K^+^/water SF configurations that occur in the two proposed permeation mechanisms, soft and direct knock-on, are shown. (*B*) Selectivity filter configurations observed in WT (PDB ID 5vk6), G77A (PDB ID 6nfu) and T75A (PDB ID 6by3) KcsA. (*C*) One possible interpretation of the fully K^+^-occupied SF observed in structures of many WT potassium channels.

In extensive experimental and computational studies of K^+^ permeation mechanisms some results support soft knock-on (e.g. interpretations of crystal structures (7, 11), streaming potentials (12–14) or global fits by electrophysiology (15)), whereas others favor direct knock-on (e.g. SF occupied by 3 to 4 K+ in anomalous diffraction data (16–18), or ‘KKKK’ configuration (Fig. 1*B*), computational electrophysiology molecular dynamics simulations (9, 10) or solid-state NMR measurements (19, 20)) (4). Thus, the debate on which mechanism actually takes place in K^+^ channels remains unresolved. Within this debate, the data on SF mutants: G77A and T75X (X=A, C, G) in the model K^+^ channel KcsA, present an intriguing case. The G77A mutation effectively removes the S3 binding site, as the backbone carbonyls of V76 point away from the ion pathway (carbonyl ‘flip’) in the crystal structure (Fig. 1*B*) (21). Interestingly, the structure shows both S3 and S1 occupied not by K^+^, but by what has been interpreted as water molecules instead, or the ‘WKWK’ configuration (however, it should be noted that without anomalous diffraction data, it’s difficult to tell whether this density corresponds to water molecules or e.g. reduced K^+^ occupancy (17)). As mentioned above, one of the arguments in favor of direct knock-on is the ‘KKKK’ configuration in crystal structures of most WT K^+^ channels. However, an alternative interpretation posits the ‘KKKK’ configuration as a superposition of soft knock-on ‘WKWK’ and ‘KWKW’ configurations (8) (Fig. 1*C*). In this context, the structure of G77A was originally interpreted as one isolating the ‘WKWK’ configuration, the same configuration that would occur during permeation in WT via the soft knock-on mechanism (21). In the same spirit, T75C and T75G mutations not only remove the S4 binding site, but lower the K^+^ occupancy at S2, which has been interpreted as an isolated ‘KWKW’ configuration (Fig. 1B) (22, 23). Interestingly, the occurrence of soft knock-on-specific configurations in G77A and T75X mutants has been taken as evidence for the soft-knock on mechanism in WT K^+^ channels (21). Functionally, mutations at both 75 and 77 positions largely decrease K^+^ conductance (ca. 32-fold for G77A (21) and between 3-17-fold for T75X (22, 24)). The data on ion selectivity are ambiguous: liposome flux assay (LFA) experiments showed WT-like K^+^/Na^+^ selectivity in KcsA G77A (21), whereas no selectivity measurements have been conducted (to the best of our knowledge) on T75X mutants. For other channels with the conserved SF sequence (TVGYG), the electrophysiological selectivity measurements are conflicting: hKv1.5 T480A and Shaker T442A (24) retained K^+^ selectivity, while NaK2K T63A and MthK T59A were non-selective (25).

In this work, we study ion permeation mechanisms in the G77A and T75A mutants of KcsA using molecular dynamics (MD) simulations with applied voltage with two modern fixed-charge force fields. In our simulations, WT KcsA permeates strictly via water-free direct knock-on, whereas both mutations drastically affect the SF conformational dynamics, coupled with permeation mechanisms incompatible with direct knock-on. In fact, the strict water-free K^+^ permeation is compromised in G77A and T75A, leading to largely reduced permeation rates, in good agreement with experimental data (21, 22, 24). The K^+^/Na^+^ selectivity of the mutants is similarly diminished. Our findings not only present a case when the structural and mutagenesis data should be interpreted with caution in the context of ion permeation mechanisms, but also support the idea that exclusion of water from the SF is necessary for the hallmark high conductance/high selectivity combination in K^+^ channels.

## Results

### Ion permeation in the WT channel

First, we simulated ion conduction in WT KcsA at the membrane voltage of 300 mV, using the CHARMM36m (26) and Amber14sb (27) force fields. In simulations with CHARMM36m we observed 40 K^+^ permeation events within the cumulative 10 µs simulation time, the majority of which occurred via direct knock-on. In 5 out of 10 simulation replicas, a water molecule entered the central parts of the SF (S2/S3). In these cases further ion permeation was halted (*SI Fig. 1A)*, suggesting that water in the SF transiently blocks ion permeation. In one simulation replica, the SF underwent drastic conformational changes with a large increase in diameter at S3, S2 and S1 that allowed permeation of both K^+^ and water (*SI Fig. 1B)*. In contrast, in Amber14sb, the SF was stable and inaccessible to water (*SI Fig. 1C*), allowing to observe 81 permeation events that occurred exclusively via direct knock-on. The calculated outward conductances are therefore 2.1±1.1 pS and 4.3±1.1 pS in CHARMM36m and Amber14sb, respectively, an order of magnitude lower than the experimental values for KcsA (3) - a known effect of K^+^ channel simulations using fixed-charge models (4).

The SF of WT KcsA in MD simulations with the CHARMM36 force field can transition toward the ‘constricted’ conformation, related to C-type inactivation, much faster than in experiments - within hundreds of nanoseconds instead of seconds (28–30). While we observed instabilities in the SF in certain simulation replicas, such transitions did not occur in our simulations (*SI Fig. 2*), possibly due to a lower temperature in our study, or limited sampling. We noticed however that the SF was on average narrower in CHARMM36m compared to Amber14sb at G77 (by 0.048±0.005 nm, distance between CA atoms of opposing subunits) and G79 (0.024±0.005 nm) (*SI Fig. 3A)*. Our previous work on MthK channels showed that subtle changes in the SF diameter can have a major effect on outward currents (31), thus it is feasible that such an effect might be present in KcsA as well.

**Fig. 2.**
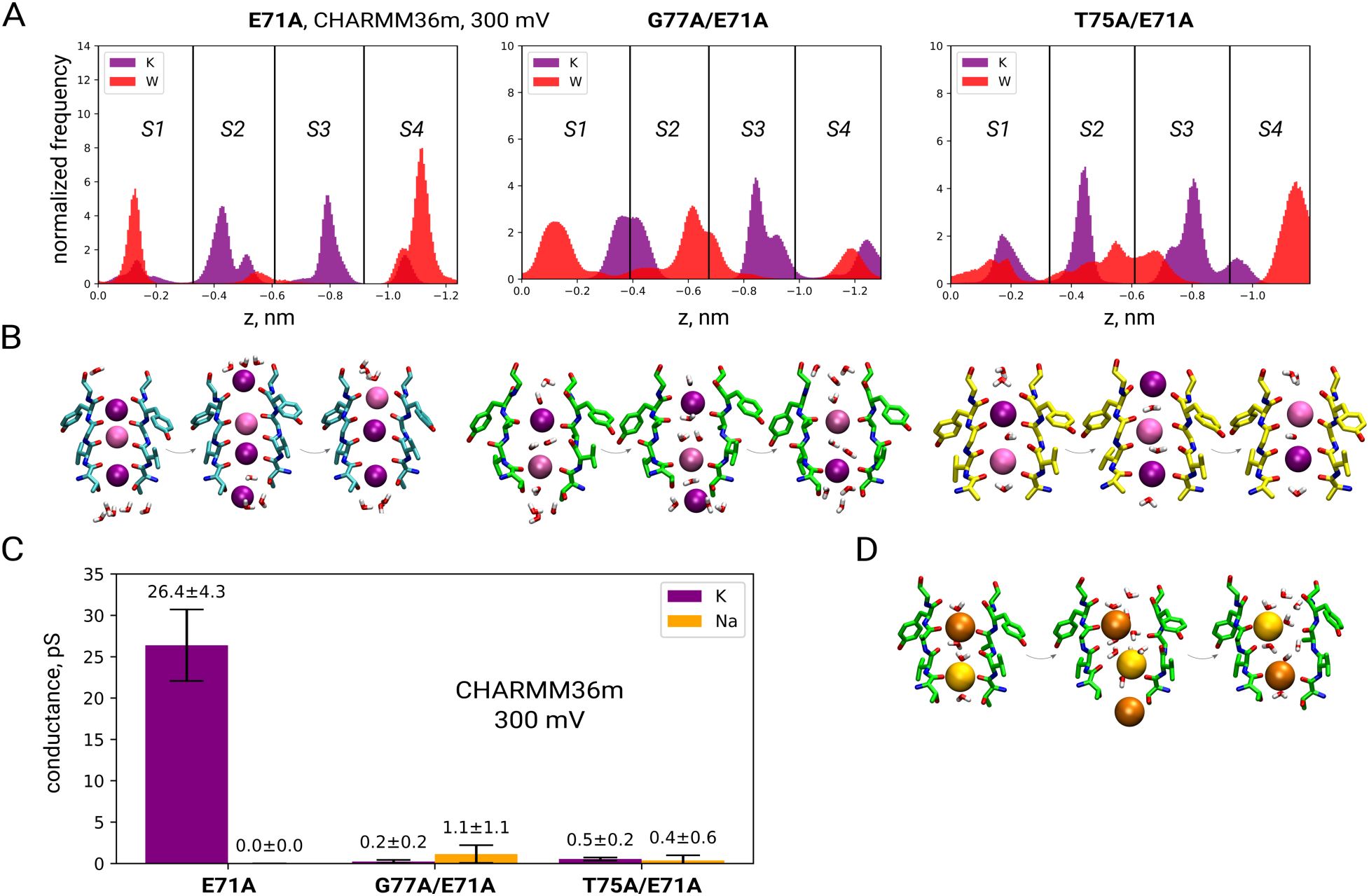
(*A*) K^+^/water distributions along the *z*-axis in SF of KcsA E71A, G77A/E71A and T75A/ E71A from all simulation replicas in CHARMM36m at 300 mV, with (*B*) respective representative K^+^/water configurations corresponding to a single K^+^ permeation event. Shift to a mechanism involving co-permeation of K^+^ and water is observed upon introduction of G77A and T75A mutations. (*C*) Mutations reduce the K^+^ conductance and impair selectivity against Na^+^ (CHARMM36m, 300mV), with (*D*) showing the representative SF configurations observed in a Na^+^ permeation event in KcsA G77A/E71A.

**Fig. 3.**
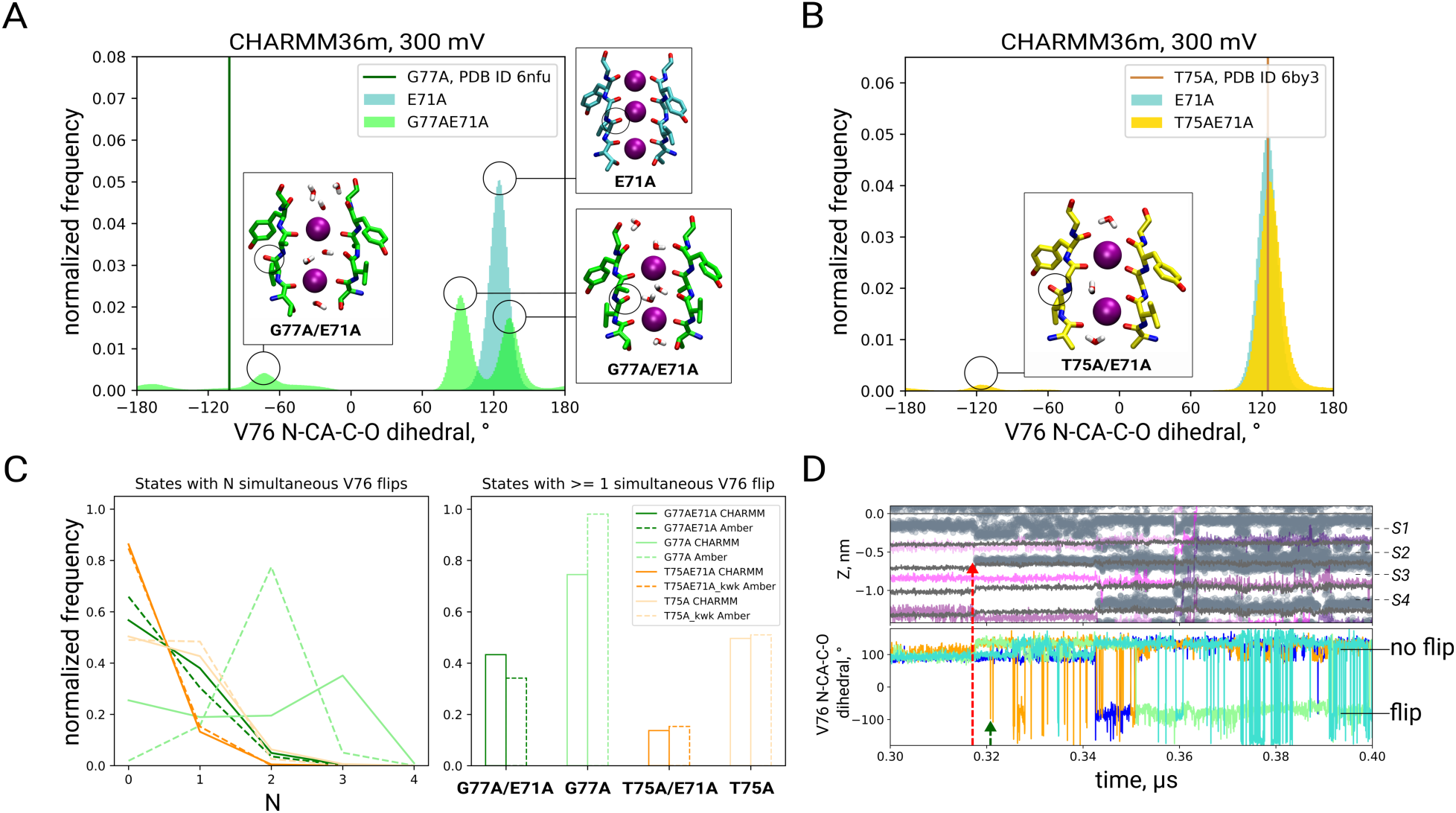
SF mutants had a higher frequency of V76 carbonyl flipping, represented by V76 N-CA-C-O dihedral angle distributions for (*A*) E71A and G77A/E71A KcsA, with the dihedral angle in the G77A KcsA crystal structure indicated (PDB ID 6nfu), and (*B*) E71A and T75A/E71A KcsA, similarly indicating the dihedral angle in the T75A KcsA crystal structure (PDB ID 6by3). (*C*) Fraction of states with various numbers (N) of simultaneously flipped carbonyls (left), and the fraction of states where at least one V76 carbonyl is flipped (right). In T75A and T75A/E71A in Amber14sb flipping has been detected only when a soft knock-on-like starting SF configuration was used (‘kwk’). (*D*) Time traces of K^+^ ions (purple lines) and water molecules (gray scatter), oxygens of SF backbone carbonyls and of threonine hydroxyls (gray lines) (up), aligned with time traces of V76 N-CA-C-O dihedral for G77A/E71A, illustrate how entrance of water into S2/ S3 (red arrow) promotes V76 carbonyl flipping (green arrow).

Finally, we tested the selectivity of WT KcsA by simulating in the presence of Na^+^ ions (instead of K^+^). To note, selectivity is often used to describe ion permeation in conditions when two ion species are present simultaneously (biionic conditions); in this manuscript however, we will use the term ‘selectivity’ for both single ion and biionic conditions. We did not observe any Na^+^ permeation in WT KcsA using CHARMM36m, and only 2 permeation events in Amber14sb (PNa^+^/PK^+^ = 0.03±0.05), showing strict K^+^ selectivity, in agreement with experiments (3).

### Ion permeation in KcsA E71A

As the WT KcsA showed unsustained K^+^ permeation frequently blocked by water, and overall low conductance, we investigated its non-inactivating E71A mutant (*SI Fig. 4A*), to obtain a clearer picture of K^+^ permeation, since earlier MD simulations as well as experiments showed higher currents and stability of KcsA E71A compared to WT (29, 32). Similarly, in our simulations, KcsA E71A showed sustained currents in both force fields. K^+^ permeation occurred consistently via direct knock-on, with rare events of water molecules entering the SF in CHARMM36m that were however able to leave so that water-free permeation continued (*SI Fig. 4B*). In total, we observed 494 permeation events over the cumulative 10 µs of simulations with KcsA E71A in CHARMM36m, and 124 in Amber14sb (*SI Table 1*) - corresponding to 26.4±4.3 pS and 6.6±1.4 pS respectively. The increased conductance of E71A correlated with the SF width: E71A is 0.05±0.02 nm wider at G79 compared to WT (CHARMM36m, *SI Fig. 3B*), in line with our previous work on MthK and its KcsA-like V55E mutant (31).

**Fig. 4.**
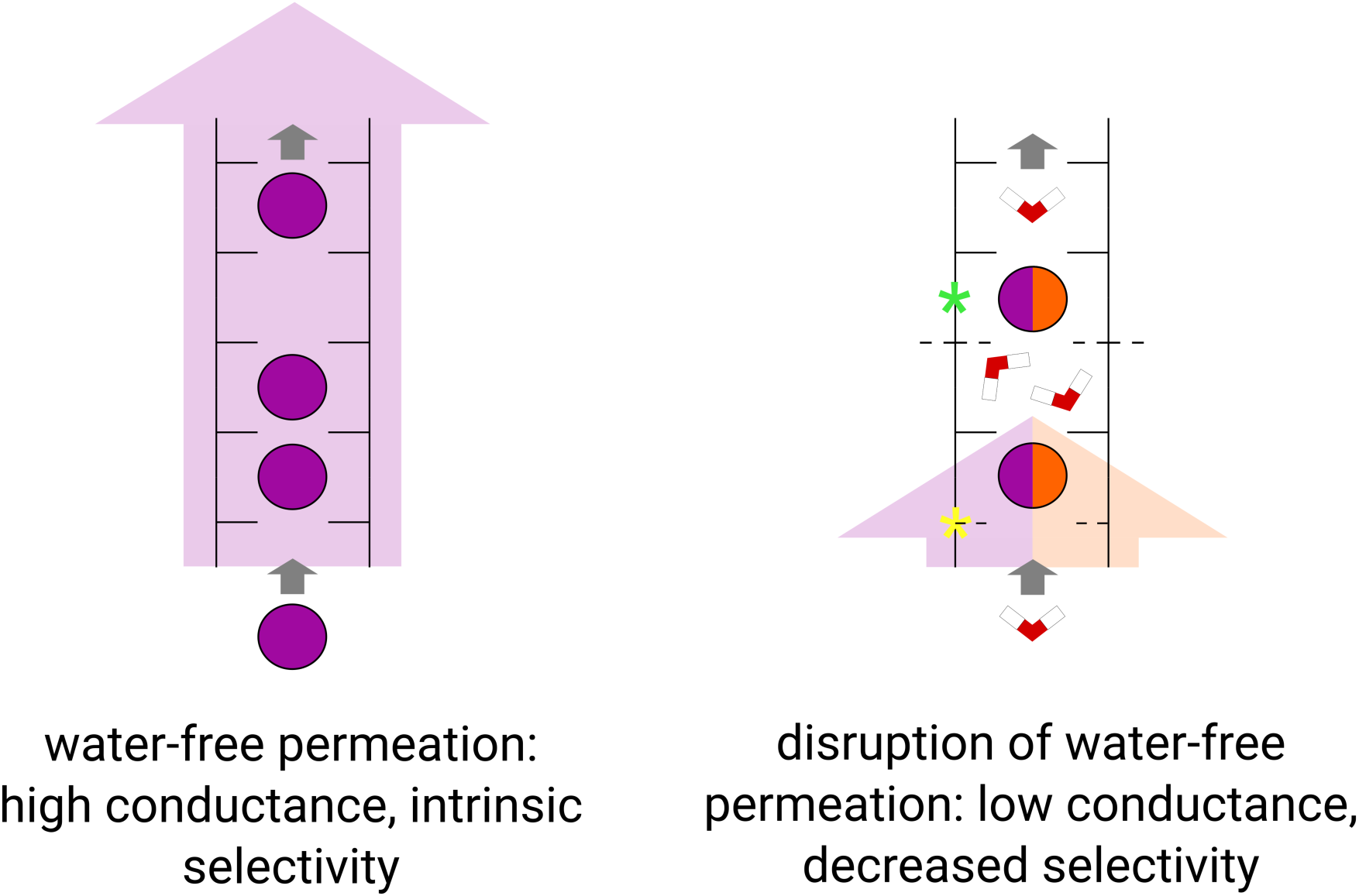
Schematic view of ion permeation in WT K^+^ channels (left) and the SF mutants (right) affecting either S4 (yellow) or S2 (green) K^+^ binding sites. While K^+^ permeation via direct knock-on is characterized by high conductance and high selectivity for K^+^ ions compared to Na^+^, disrupting the water-free permeation by e.g. introducing mutations to the SF impedes both permeation rates and ion selectivity. Both mutations we studied promoted flipping of V76 carbonyls (right, dashed lines in the middle), regardless of whether they affected the adjacent G77 or distant T75, which we also link to the destabilizing effect of water presence in the SF.

At the same time, KcsA E71A retained the strict K^+^ selectivity: no Na^+^ permeation was observed in simulations with the CHARMM36m force field, and only 3 Na^+^ permeation events in Amber14sb (PNa^+^/PK^+^ = 0.02±0.04). It should be noted that the experimental measurements of E71A K^+^/Na^+^ selectivity are not fully clear: while ^22^Na^+^ flux measurements of E71A suggested lower selectivity compared to WT (33), reversal potential shifts showed no significant change in selectivity in E71A (33, 34). A possible explanation for this observation is the propensity of the E71A SF to adopt a specific conformation in pure Na^+^ conditions (PDB ID 3OGC (33), although a similar conformation has been observed in K^+^-containing solution as well, PDB ID 2ATK (34)), not seen in WT KcsA (as it inactivates instead). This E71A-specific conformation shows flipped V76 carbonyls and flipped side chains of the residue D80. In fact, we observed rare hints of such conformation in our simulations of E71A in pure Na^+^ (*SI Fig. 5*), although likely due to the limited sampling and therefore short simulated times with this conformation, we did not record any Na^+^ permeation events in it. Therefore, we believe we can safely assume that, for this study, WT KcsA and E71A KcsA display similar K^+^/Na^+^ selectivity, as long as their SFs retain the canonical, conductive conformation.

To summarize, in our simulations KcsA E71A showed essentially the same permeation mechanism (direct knock-on) and ion selectivity as the WT channel, albeit with much higher currents. We thus conducted our study in the following manner: first, we introduced the SF mutations into KcsA E71A, and investigated their effect on K^+^ permeation in G77A/E71A and T75A/E71A constructs. As control systems, we simulated the single G77A and T75A mutants, and compared their behavior to the ‘pure’ WT KcsA (that is, without the E71A mutation), to account for possible biases of channel sequences and force fields, and get the comprehensive picture of the effect of SF mutations on ion permeation.

### Effect of the G77A mutation

The introduction of G77A into KcsA E71A led to a drastic change in the permeation mechanism: the mutated channel co-permeated water molecules, thus compromising the water-free direct knock-on mechanism (*Fig. 2A, B, middle*). Although this mechanism resembled soft knock-on, there were some differences - K^+^ ions permeated by jumping directly from the KW**K**W configuration to **K**WKW - without the intermediate W**K**WK that is expected according to the canonical definition of the soft knock-on mechanism (*SI Fig. 6A*). Additionally, the number of intervening water molecules between K^+^ ions was often 2 or more (instead of expected 1 water molecule). The dominance of the KWKW occupancy pattern is not compatible with electron densities in the SF of WT channels that correspond to similar ion occupancy for all four central ion binding sites (6, 11, 17), hence rendering this permeation mechanism unlikely for the WT channel. Importantly, the average position of K^+^ and water inside the SF in our simulations is in good agreement with the crystal structure of G77A (21) (*Fig. 2A*), with an exception of S4. In the crystal structure, S4 is occupied by a K^+^ ion, whereas simulations show a larger K^+^ density at S3 in both force fields (*Fig. 2A* and *SI Fig. 7A*). This discrepancy may be related to V76 carbonyl flipping dynamics: in the crystal structure, all 4 V76 are flipped, while in our simulations the number of simultaneously flipped V76 ranged between 0 and 4, and conformations with a larger number of simultaneous flips were less frequent (*Fig. 3C*). Accordingly, the fraction of frames where K^+^ occupied S4 instead of S3 increased with the number of simultaneously flipped V76 (*SI Fig. 7B*).

The above mentioned shift in the ion permeation mechanism was accompanied by a drastic reduction in K^+^ conductance for G77A/E71A compared to E71A. In total, we observed only 16 K^+^ permeation events in CHARMM36m, and 4 in Amber14sb, over ∼30 µs and 10 µs total simulation time, respectively (*SI Table 1*), corresponding to an average ∼100-fold (CHARMM36m) and ∼30-fold (Amber14sb) reduction in outward conductance (*Fig. 2C* and *SI Fig. 8A*) due to the mutation. This decrease agrees well with experimental estimates (32-fold reduction) (21). Further, K^+^/Na+ ion selectivity of G77A/E71A was clearly impaired in CHARMM36m, as the channel permeated Na^+^ ions at a rate similar to K^+^ (*Fig. 2C*). Na^+^ permeation, akin to K^+^, featured water co-permeation (*Fig. 2D, SI Fig. 6B*). While the loss of ion selectivity has a straightforward explanation within the direct knock-on framework (10) (lack of complete ion desolvation reduces K^+^ vs Na^+^ selectivity, due to differences in solvation free energies), it is in apparent contrast with the interpretation of the LFA experiments in which the G77A mutant does show ion selectivity (21). We address these discrepancies in the Discussion. To get a more complete comparison to the LFA measurements, where the channel is oriented randomly in the liposomal membrane, we also simulated inward permeation (at −300 mV) in CHARMM36m. Similarly to simulations at positive voltage, G77A/E71A was equally permeable to Na^+^ and K^+^, while E71A retained its selectivity (*SI Fig. 8B*). In contrast, we did not observe any Na^+^ permeation in simulations in Amber14sb. Given the very low K^+^ permeation rate of G77A/E71A with this force field (4 events in 10 µs), it is difficult to conclude whether this mutant is actually selective in this force field. Finally, we tested biionic conditions with two K^+^/Na^+^ ratios (2:1 and 1:2) (*Table 1*); however, we saw little to no permeation of either ion in both CHARMM36m and Amber14sb at +300 mV, and some permeation of both ions in CHARMM36m at −300 mV (*SI Fig. 8C*). Control simulations of KcsA G77A without the E71A mutation also showed a water-mediated permeation mechanism, with a reduced average outward conductance compared to WT in CHARMM36m (∼2-fold) (*SI Fig. 9A*). Ion selectivity was compromised in CHARMM36m at both positive and negative voltages, as well as in biionic conditions (tested at +300 mV) (*SI Fig. 9B-D*). In Amber14sb, we did not observe K^+^ or Na^+^ permeation, whereas flipping of V76 occurred in both force fields (*Fig. 3C*).

### Effect of the T75A mutation

T75A removes the hydroxyl groups that form the bottom of the S4 ion binding site. Consequently, in the crystal structure of KcsA T75A the K^+^ ion in S4 is replaced by a water molecule (24) (*Fig. 1B, right*). In our simulations of KcsA T75A/E71A, this had a dramatic effect on K^+^ permeation, in a force field-dependent manner. In Amber14sb we did not observe any ion permeation, regardless of whether the initial SF configuration was direct or soft knock-on-like (*SI Table 1*). In contrast, in CHARMM36m, the introduction of the T75A mutation led to co-permeation of K^+^ and water (*Fig. 2A, right, SI Fig. 10*). Interestingly, while in the crystal structure of KcsA T75A the SF carbonyls are not flipped, we observed a tendency of T75A/E71A to flip at V76, in a similar manner to G77A/E71A and G77A (*Fig. 3A-C).* Flipping correlated with water entry into the SF, suggesting that it could be a general feature of water-filled selectivity filters (*Fig. 3D, SI Fig. 12C*).

KcsA T75A/E71A in CHARMM36m showed a low outward K^+^ conductance of 0.5±0.2 pS (*Fig. 2C*) (10 permeation events in 10 µs), which is in good agreement with experiments (24). Further, the mutant was equally permeable to K^+^ and Na^+^ (*Fig. 2C*), suggesting compromised ion selectivity. We cannot compare this prediction with experiments, as ion selectivity measurements are lacking for KcsA T75X constructs, and the data for other K^+^ channels are inconclusive (see Introduction).

In simulations of KcsA T75A without the E71A mutation, we also observed low K^+^ conductance of 0.40±0.16 pS (15 K^+^ permeation events over 20 µs) in CHARMM36m (*SI Fig. 11A*), and no ion permeation in Amber14sb. The permeation mechanism was generally similar to that of T75A/E71A, featuring co-permeation of water and K^+^. Interestingly, while the double mutant showed a preference for a stepwise motion of the KWK pair through the SF (*SI Fig. 10*), here, direct jumps from KW**K**W to **K**WKW occurred, without the intermediate configuration W**K**WK (*SI Fig. 11C*). Again, this is at odds with the original soft knock-on mechanism that requires KWKW and WKWK to occur with similar probabilities, in order to explain the similar electron densities at the four central ion binding sites in the SF of non-mutated channels. Accordingly, the resulting K^+^ density profile displays a relatively diminished K^+^ density at S2 compared to the double mutant (*SI Fig. 15*), reminiscent of the crystal structure of KcsA T75C and T75G (22, 23). The frequency of V76 flipping was higher compared to the double mutant *(Fig. 3C*), possibly explaining the reduced K^+^ density at S2 by perturbed K^+^-V76 carbonyl interactions. Regarding ion selectivity, 2 Na^+^ permeation events in single ion conditions over the total 25 µs of simulation were detected (conductance of 0.05±0.08 pS).

As noted before, the starting structures for all simulations mentioned were prepared by introducing the appropriate mutations into the KcsA E71A crystal structure. However, a crystal structure of KcsA T75A in the open state is available (24). We thus built and performed simulations of this KcsA T75A structure, together with WT KcsA back-mutated from the same structure (CHARMM36m, +300 mV). The results were in broad accord with our main simulation set, with permeation via direct knock-on in WT KcsA, with occasional water occupancy in the SF, low conductance but K^+^/Na^+^ selectivity, and soft knock-on with low conductance in T75A. Interestingly, while the preference for **K**WKW to KW**K**W jumps wasn’t as pronounced compared to T75A from the main simulation set, the relative K^+^ density drop in S2 was nevertheless present (*SI Fig. 16*). Additionally, in this new system we could detect a higher number of both K^+^ and Na^+^ permeation events, and conclude lower selectivity in T75A with higher confidence (*SI Fig. 11B*).

## Discussion

In this work, we studied the effect of the G77A and T75A mutations on the ion permeation mechanism and selectivity in the model potassium channel KcsA channel using MD simulations with applied voltage, using two popular force fields. We show that the introduction of these mutations leads to a complete change of the K^+^ permeation mechanism - from a water-free direct knock-on to mechanisms that feature co-permeation of water. This, in turn, dramatically reduces ion permeation rates, in good agreement with experimental data. Importantly, we introduced mutations into the SFs of both WT KcsA and its non-inactivating E71A variant; this suggests that the observation of the shift towards water-mediated permeation mechanisms and low conductance is solely due to the introduced mutations and independent of the initial channel.

The shift in the permeation mechanism is rationalized by the altered conformational landscape of mutated SFs, reminiscing our previous work on selective and non-selective permeation in K^+^ channels and their non-selective counterparts, as well as single molecule FRET experiments from the Nichols lab. Indeed, we previously observed dilated and more dynamic SFs of non-selective channels that allow simultaneous permeation of ions and water molecules, and we linked this increased water co-permeation to reduced K^+^/Na^+^ selectivity (10). Of particular interest, a series of FRET experiments on KirBac1.1 (35), NaK2K and TREK-2 (36) channels showed similar, ion-dependent SF conformational states in all of these channels. In these experiments, the presence of K^+^ in the SF promotes a compact, Rb^+^-permeable conformation of the SF, whereas in Na^+^ the SF adopts a dilated, Na^+^ and water permeable state, thus offering an explanation how water-free direct knock-on can occur via compact states, and yet still display water permeation through Na^+^-induced dilated conformations. Another study on S3 glycine mutants in KirBac1.1 also showed more dynamic SFs compared to WT and consequently reduced K^+^/Na^+^ selectivity (37). In simulations presented in this work, the main source of increased SF dynamics is associated with V76 carbonyls. In the crystal structure of KcsA G77A, S2-and S3-forming V76 carbonyls are in the flipped state as they point away from the pore axis (21) (*Fig. 1B, middle*). This V76 flipping occurs spontaneously in our simulations of G77A/E71A (and G77A). In contrast to the crystal structure however, not all 4 V76 are simultaneously flipped throughout the simulations, but rather the number of flipped carbonyls fluctuates between 0 and 4. The details of carbonyl dynamics vary between the systems (with or without the E71A mutation) and the force field used (*Fig. 3C*). These differences may be explained by the fact that the crystal structure was solved with an imposed four-fold symmetry and also at a lower temperature (100 K) than the one we used in our simulations (290 K). Consequently, low energetic barriers between flipped/non-flipped states are crossed. Notably, this carbonyl dynamics directly affects the position of the lower ion in the SF, as it is more often located in S4 (i.e. its crystallographic site) when more V76 are simultaneously flipped (*SI Fig. 7B*).

Unexpectedly, we observed a similar behavior in the T75A mutant. Even though it only removes the hydroxyl groups of the S4-forming T75 residue, and otherwise does not affect the SF conformation in the crystal structure (*Fig. 1B, right*) (24), we observed flipping of V76 carbonyls in simulations of T75A as well. A unifying factor was a switch (either transient or not) to a water-mediated permeation mechanism; indeed, for all systems we studied, we found a clear correlation between V76 flipping and water entrance into the K^+^ binding sites formed by V76, S2 and S3, including WT KcsA and E71A that showed some frequency of flipping too (*SI Fig. 12A, B*). Similar correlation between water presence in the SF and flipping of S2/S3 carbonyls was demonstrated in previous MD studies of MthK (38), TREK-2 (39) and HERG (40) channels. We analyzed the causal relationship of flips and other factors; the vast majority of flipping events happened when water was already in S2/S3 for most systems, despite a sizable fraction of frames (CHARMM36m) with no water in S2/S3, thus further supporting the preference of flipped states for water presence in the SF (*SI Fig. 12C*). G77A in CHARMM36m was a notable exception, with more than 30% of flips starting with no water in S2/S3, indicating also the role of intrinsic propensity to flip of a given channel. Interestingly however, flipping events that started with water in S2/S3 tended to last longer in all systems (*SI Fig. 12D*). While we previously described the ion occupancy of lower sites (S3/S4) correlating with the number of simultaneous V76 flips, we did not find the absence of ions in S2/S3 to be necessary for flips to occur (*SI Fig. 13*).

A plausible structural explanation for the effect of water on flips could be in the several types of hydrogen bond networks that water molecules formed with carbonyl oxygens. First, water interacted with V76 carbonyls while inside the SF, which could potentially stabilize some and destabilize other flipping configurations (*SI Fig. 14A*). Second, water behind the SF (primarily in G77A/E71A and T75A/E71A, where bulky glutamates behind the SF were removed) could sometimes interact with and stabilize the flipped V76 (*SI Fig. 14B*). Taken together however, carbonyl flipping is likely affected by a combination of factors, such as the presence and configuration of water molecules in and around the SF, potentially ion configuration, as well as the intrinsic propensity to flip of specific SF sequences and the regions behind it (*Fig. 3C*).

The exquisite ion selectivity of K^+^ channels arises from a balance of interactions between ions, water and the SF. We have previously shown that the direct knock-on mechanism is intrinsically selective for K^+^ against Na^+^, due the difference in ion dehydration free energies (10), therefore we expected a diminished ion selectivity for the water permeating mutants with dilated SFs. In line with these expectations, mutants of other K^+^ channels at the second glycine in the SF (corresponding to G77 in KcsA), i.e. BK G354S (41), GIRK2 G156S (human G154S) (42, 43) and Shaker G376A (5), show a loss of ion selectivity in electrophysiology. On the contrary, LFA measurements of KcsA G77A suggested that it conserves ion selectivity, and isothermal titration calorimetry showed similar binding affinities of ions to the WT and mutant channels (21). In our simulations both G77A and G77A/E71A are clearly non-selective in CHARMM36m, at both positive and negative voltages. G77A in CHARMM36m, where we were able to detect ion permeation when both K^+^ and Na^+^ were present, was non-selective in these conditions as well. In Amber14sb these mutants did not display any Na^+^ permeation, but also K^+^ conductance was very low (i.e. only 4 permeation events in 10 µs for G77AE71A, and G77A was non-conductive), which prevented us from making conclusions about their ion selectivity with this force field. One possible reason for this discrepancy between LFA experiments and MD simulations could be the different driving forces of ion permeation: while an applied electric potential was used in our study, in LFA ions are driven in or out of liposomes by their concentration gradients. This can lead to uncertainty in the instantaneous transmembrane voltage. Second, the intracellular cavity of KcsA was shown to be blocked by Na^+^ in a voltage-dependent manner (44). Na^+^ block can be relieved either by diffusion of Na^+^ back into the intracellular solution, or via a ‘punchthrough’ mechanism through the SF by incoming ions; however, it is not clear to what extent this phenomenon is present for KcsA in LFA and how it would affect the results. Then, the method showed an overall higher current for K^+^ compared to Na^+^ in this setup, but did not preclude Na^+^ currents, nor quantify the effect of G77A on ion selectivity. In this vein, a possible interpretation of the LFA results that could be reconciled with our MD data is that even if KcsA G77A possesses some preference for K^+^ against Na^+^, the selectivity is still diminished compared to WT, as it is the case for the mutants of the glycine of BK (PNa^+^/PK^+^ ∼ 0.2) (41), GIRK (∼0.8) (42, 43) and Shaker (∼0.8) (5) channels. Finally, it is also not straightforward to interpret the similar binding affinities in ITC, as it does not provide energetic barriers crucial for ion permeability, which is a kinetic property.

Notably, mutations of the S3 glycine in K^+^ channels in humans are associated with disorders such as progressive cerebellar ataxia (BK G354S (41)) and Keppen-Lubinsky syndrome (GIRK2 G154S (45)). BK and GIRK2 are expressed in multiple tissues in humans, and participate in many processes including control of the smooth muscle tone and contributing to the resting potential of neurons, respectively (46, 47). As mentioned above, these mutants have diminished conductance and ion selectivity, and in a previous MD study of GIRK2 G154S water-mediated K^+^ permeation was reported (48). Taken together with our results, it is plausible that a shift in the ion permeation mechanism happens in the mutants of these channels as well, disrupting proper ion permeation and leading to these disease phenotypes. This could provide a mechanistic basis that may guide future development of drugs targeting those disorders.

The T75A mutation behaved similar to G77A in our simulations: it displayed reduced selectivity for K^+^ (for both single and double mutations) in CHARMM36m at positive voltage (we did not observe any K^+^ or Na^+^ permeation for either system at negative voltage or in Amber14sb), with identical K^+^ and Na^+^ conductances in T75A/E71A, and somewhat higher conductance for K^+^ in T75A (*Fig. 2C, SI Fig. 11A, B*). As mentioned, we are not aware of ion selectivity measurements for KcsA T75A. However, the ion selectivity measurements of other K^+^ channels with mutated threonine at the same location are conflicting: Kv1.5 T480A and Shaker T442A mutants were selective (24), while NaK2K T63A and MthK T59A were non-selective (25). This channel-dependent effect of the SF threonine substitution can indicate channel-dependent shifts in the ion permeation mechanism, posing an intriguing question - do selective mutants conserve the WT permeation mechanism, while non-selective ones have it switched to a non-WT mechanism, similarly to what we observed for KcsA T75A?

Recently, the fact that crystal structures of G77A and T75X mutations of KcsA show ion/water configurations of their SFs has been interpreted as evidence for the soft knock-on mechanism in WT channels as well (21). In this context, the ‘KKKK’ configuration found in SFs of WT channels is viewed as a superposition of ‘WKWK’ and ‘KWKW’ configurations (characteristic for soft knock-on, *Fig. 1A*). Consequently, KcsA G77A has been proposed to isolate ‘WKWK’, whereas KcsA T75X would isolate ‘KWKW’, as T75X showed a decreased K^+^ density at S2 (on top of eliminating S4). This presents an argument in favor of soft knock-on in WT KcsA, under a critical assumption that the permeation mechanism is not affected by the introduced mutations. Our results, however, show that both mutations largely perturb the conformational landscape of the SF, to an extent that makes the permeation mechanism in the mutants incompatible with that of the WT. Thus, we argue the experimental data on KcsA G77A and T75X do not contradict the direct knock-on mechanism in WT K^+^ channels. Instead, G77A and T75X simply conduct ions (and water) via an entirely different mechanism. Water-free K^+^ permeation within the direct knock-on framework as a prerequisite for combined high conductance and selectivity of K^+^ channels is supported by several studies, both computational and experimental (4), and our current results fit very well within this framework, thus further solidifying direct knock-on as the permeation mechanism in K^+^ channels.

## Materials and Methods

### System preparation

The starting structure for the KcsA non-inactivating mutant E71A in the open conformation (PDB ID: 5vk6 (49)), embedded in a lipid bilayer, was built using CHARMM-GUI (50, 51). Its sequence (residues 26 to 121, preserving the E71A mutation) was mutated to match UniProtKB entry P0A334. N- and C-termini were acetylated and methylamidated, respectively. The system contained 39 1,2-dioleoyl-sn-glycero-3-phospho-(1’-rac-glycerol) (DOPG) and 105 1,2-dioleoyl-sn-glycero-3-phosphoethanolamine (DOPE) molecules in the lipid bilayer; 11257 water molecules; 163 K^+^ and 140 Cl^-^ ions to yield a neutral system and a K^+^ concentration of 0.8 M in the water phase. 4 of DOPG molecules were placed in their respective crystallographic binding sites in KcsA. E118 and E120 were protonated to stabilize the open state of the channel (52).

The KcsA T75A/E71A system was built by manually introducing a corresponding mutation to the KcsA E71A system (4x KcsA monomers). To obtain the WT structure, A71 in the KcsA E71A system were reverted to a glutamate which was then protonated in line with ssNMR results (53). As the KcsA E71A structure has 2 water molecules per monomer that establish the hydrogen bonding network behind the SF, compared to 1 in WT (6), the additional water molecules were removed accordingly. SF mutants without E71A were obtained by manually introducing G77A or T75A to this WT structure. KcsA G77A/E71A was built separately in CHARMM-GUI from E71A 5vk6 using a similar protocol, and contained in addition to the protein 33 DOPG and 87 DOPE molecules, 9315 water molecules, 134 K^+^ and 117 Cl^-^, resulting in an electrically neutral system with the 0.8 M concentration of K^+^ in the water phase.

Additionally, we built a system with the crystal structure of KcsA T75A in the open state (PDB ID 6by3, (24)), and a control system with this structure back-mutated to WT KcsA. Briefly, we introduced all mutations outside the SF necessary to match the sequence of other systems we simulated, and built the two systems in CHARMM-GUI. Each of the final systems contained 39 DOPG, 105 DOPE, 7700 water molecules and 114 K^+^ (0.8 M) and 91 Cl^-^ for an electrically neutral system.

After building a system in CHARMM-GUI, it was equilibrated using the default 6-step protocol and CHARMM36m/TIP3P parameters (26, 54–56) provided by CHARMM-GUI to reach the temperature of 290 K and the pressure of 1 bar. After introducing necessary mutations, the systems were additionally equilibrated for up to 100 ns in the NPT ensemble at respective temperatures and pressure (in this and all the subsequent simulations kept constant by the semi-isotropic barostat Parrinello-Rahman barostat (57), and velocity rescale (v-rescale) thermostat (58)), without restraints. Periodic boundary conditions (xyz) were used. Constraints were used for bonds with H atoms (LINCS (59)). Simulation timestep was 2 fs. Lennard-Jones interactions were treated with a cutoff (forces switched to 0 from 1.0 nm to 1.2 nm in CHARMM36m, plain cutoff at 0.9 nm with long range dispersion corrections for energy and pressure in Amber14sb). Coulomb interactions were treated with PME (60) with a 1.2 nm cutoff in CHARMM36m and 0.9 nm in Amber14sb. All simulations were carried out using GROMACS 2020.6 or GROMACS 2020.7 (61).

Amber14sb/TIP3P systems with LIPID17 lipids (27, 54, 62–64) were generated from the CHARMM36m structure immediately after the initial 6-step equilibration, using the charmmlipid2amber (63, 64), GROMACS pdb2gmx, or CHARMM-GUI force field conversion tools (65). Systems in Amber14sb were then additionally equilibrated for up to 100 ns in the NPT ensemble before turning on the voltage, similar to the CHARMM36m systems.

Systems with Na^+^ as the permeating ion were prepared by either a) replacing K^+^ with Na^+^, or b) with specific K^+^/Na^+^ ratios in select systems to simulate biionic conditions in the structures after equilibration.

### Molecular dynamics simulations with applied voltage

For production runs, a constant electric field was applied to the equilibrated structures along the *z*-axis (perpendicular to the membrane plane) to yield approximately 300 mV (or −300 mV for control simulations under negative voltage). Previously it was shown that to obtain a correct transmembrane voltage in MD simulations, the whole box size in *z* should be taken into account (66). Correspondingly, the electric field *E* was calculated using a formula *E = ΔV / dz,* where *dz* is the size of the box along the *z-*axis. For each system, 10 simulation replicas were carried out with 1-2 µs per replica. For certain mutant systems we performed control simulations starting from both direct and soft knock-on-like SF configurations. To check whether the channel stayed in the open state, we monitored the stability of the channel’s activation gate by calculating the distances between T112 CA atoms of opposing subunits; the gate remained open for the majority of simulation time (*SI Fig. 17*). All simulations we carried out are listed in *SI Table 1*.

### Data analysis

To count permeation events, a custom Python script was used, kindly provided by Vytautas Gapsys (available upon request). The simulation box was divided into 4 regions - below the SF (‘1‘), in the SF but below its midpoint (‘2‘), in the SF above its midpoint (‘3‘), and above the SF (‘4‘). A permeation event was defined as a sequential transition 1→2→3→4 for outward permeation, and in the opposite direction for inward permeation (i.e. ions that have been initially inside the SF are not counted, unless they reentered the SF later and then permeation occurred according to our definition). To calculate outward conductance, the difference between the number of outward and inward permeation events was used; accordingly, the inward conductance for systems at negative voltage was taken as the negative of that value. Presented conductance values are averages over all simulation replicas for a given system, with error bars representing 95% confidence intervals calculated using Student’s *t-*distribution.

To collect ion/water positions along the *z-*axis in SF, a custom FORTRAN code was used, available in (38). Dihedral angles were collected using the standard GROMACS toolbox. A flipped state of V76 was defined as the N-CA-C-O dihedral angle falling between −130° and −50°. For plotting the data, Python 3, matplotlib and numpy were used (67, 68). Molecular visualizations were rendered using VMD (69).

## Acknowledgements and funding sources

The authors thank Sergio Pérez-Conesa and Lucie Delemotte for the initial structures of KcsA E71A, and Vytautas Gapsys for the ion permeation counting script. The authors acknowledge funding from the Deutsche Forschungsgemeinschaft (DFG, German Research Foundation) through FOR2518 ‘Dynion’ Project P5.

## SI Figures

**SI Fig. 1.**
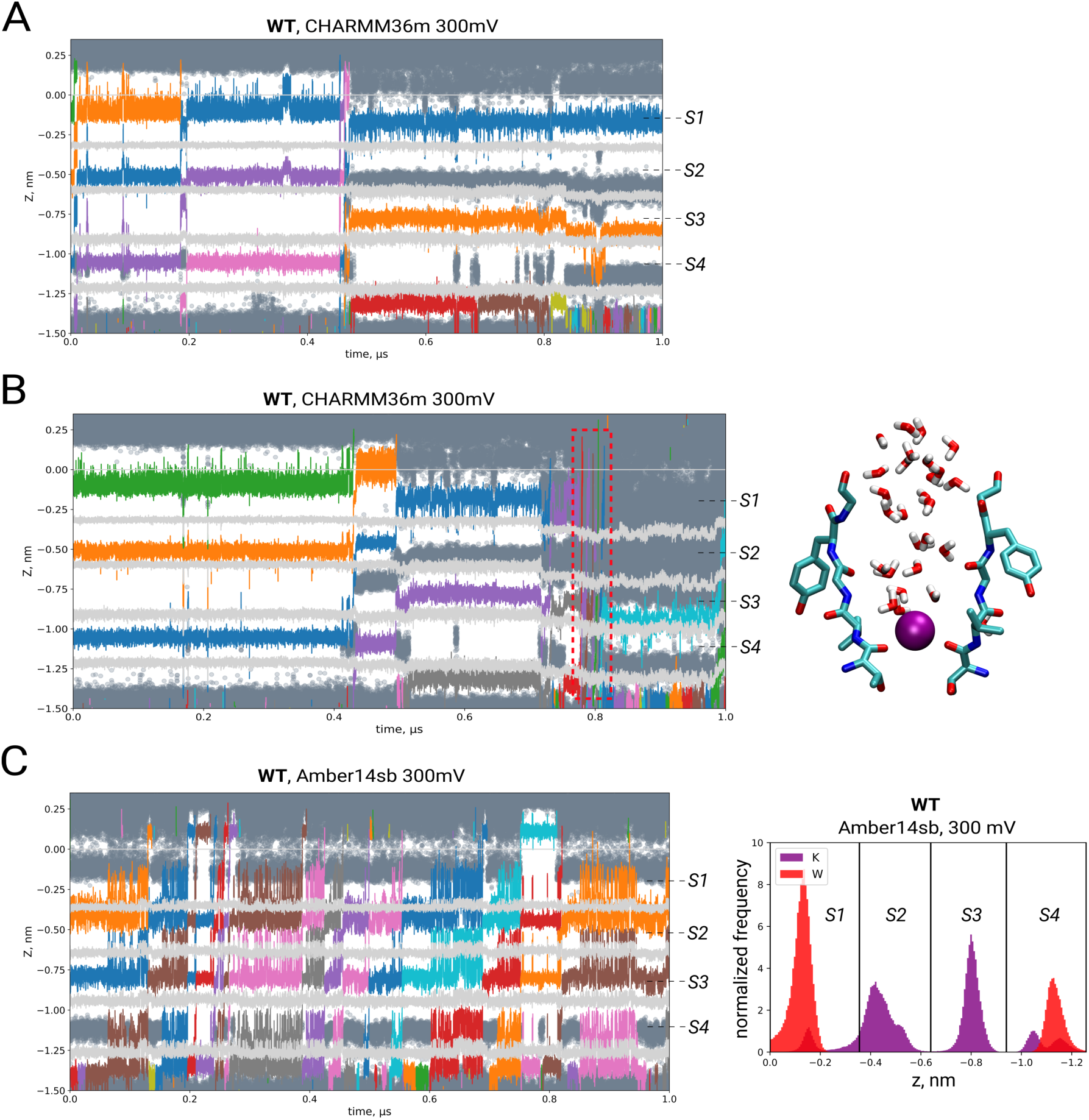
(*A*) Example traces of K^+^ ions (colored lines), water molecules (dark gray scatter) and backbone carbonyl oxygens/threonine hydroxyls (light gray lines) of the SF of WT KcsA in CHARMM36m showing K^+^ permeation via direct knock-on followed by entrance of water into the SF and block of further K^+^ permeation. (*B*) Traces showing several events of K^+^/water co-permeation in one of the simulation replicas of WT KcsA in CHARMM36m (left). Those events were accompanied by a largely distorted SF with widening in the upper regions (right). (*C*) Traces showing K^+^ permeation in WT in Amber14sb (left). Permeation occurred strictly via direct knock-on with water being virtually absent in the central regions of the SF, as demonstrated by the K^+^/water distribution from all simulation replicas (right).

**SI Fig. 2.**
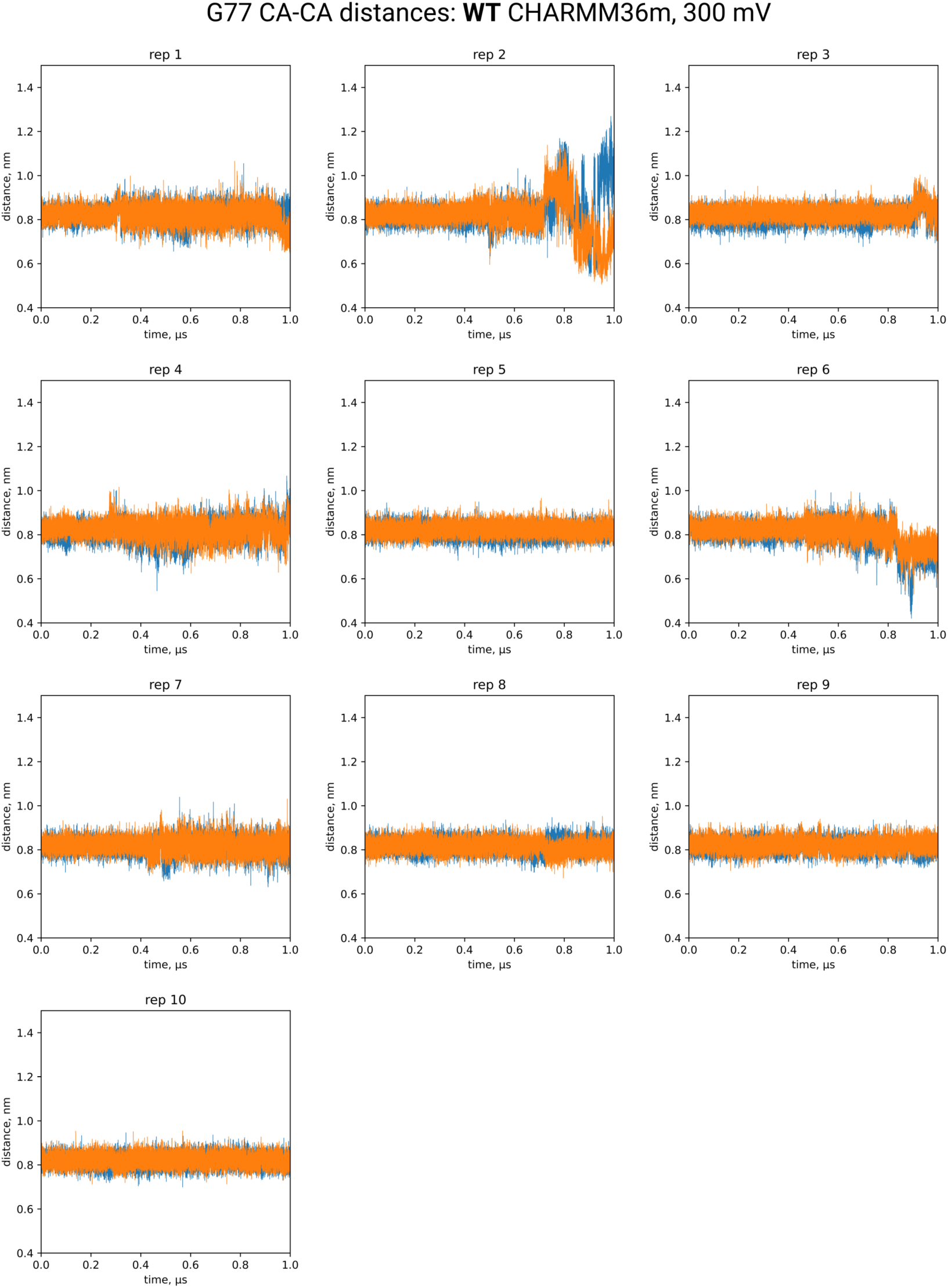
Distance between ɑ-carbons of G77 (opposing subunits) in the SF of WT KcsA. Transition to the inactivated state in WT KcsA is characterized by narrowing of the SF at G77 from ∼0.8 nm to ∼0.5 nm; in our simulations the WT SF was generally stable against this transition.

**SI Fig. 3.**
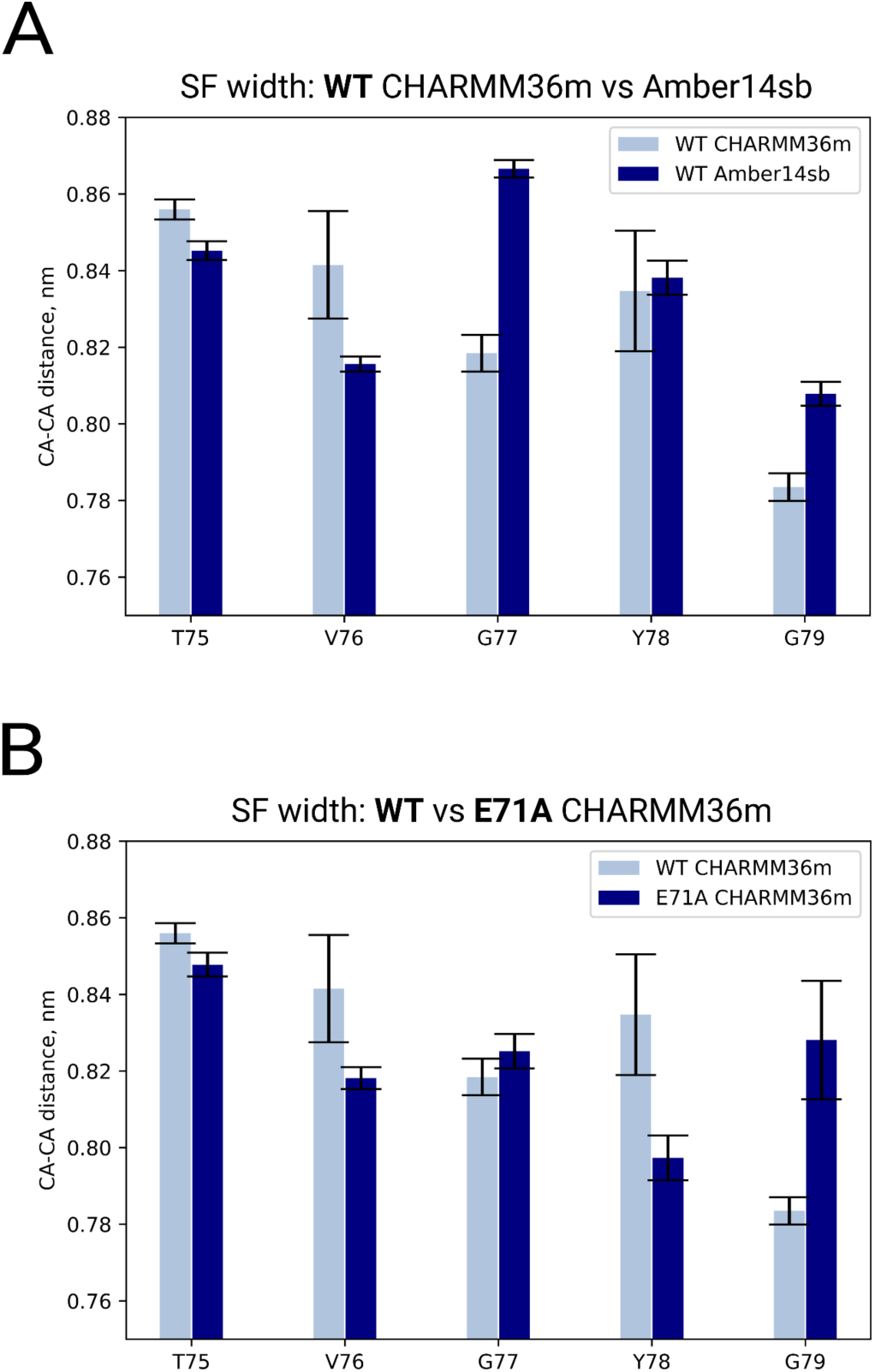
Average distance between ɑ-carbons of opposing subunits in the SF of WT KcsA in CHARMM36m and Amber14sb. Mean distances were calculated for each of 10 replicas, then the averages of these means were calculated; error bars represent CI 95% of those averages.

**SI Fig. 4.**
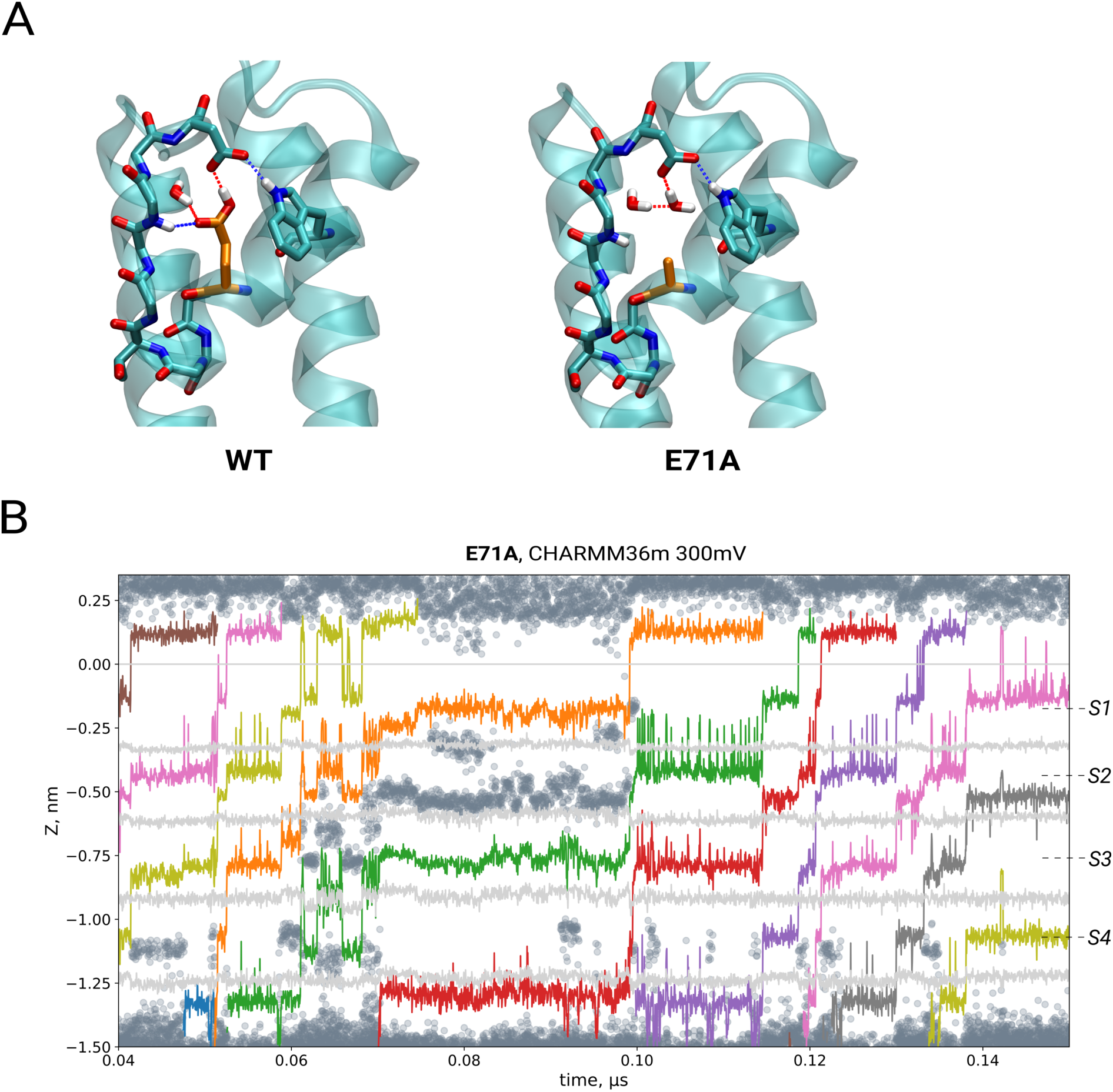
(*A*) Hydrogen bond network behind the SF of WT KcsA (left, PDB ID 1k4c (6)) essential for SF inactivation. Mutation of E71 (orange) to alanine disrupts this hydrogen bond network (right, PDB ID 5vk6 (49)) and prevents the transition to the inactivated state. (*B*) Traces of K^+^ ions (colored lines), water molecules (dark gray scatter) and backbone carbonyl oxygens/ threonine hydroxyls (light gray lines) of the SF in one of the simulation replicas of E71A in CHARMM36m show transient entrance of water into the SF, followed by continuation of ion permeation via water-free direct knock-on.

**SI Fig. 5.**
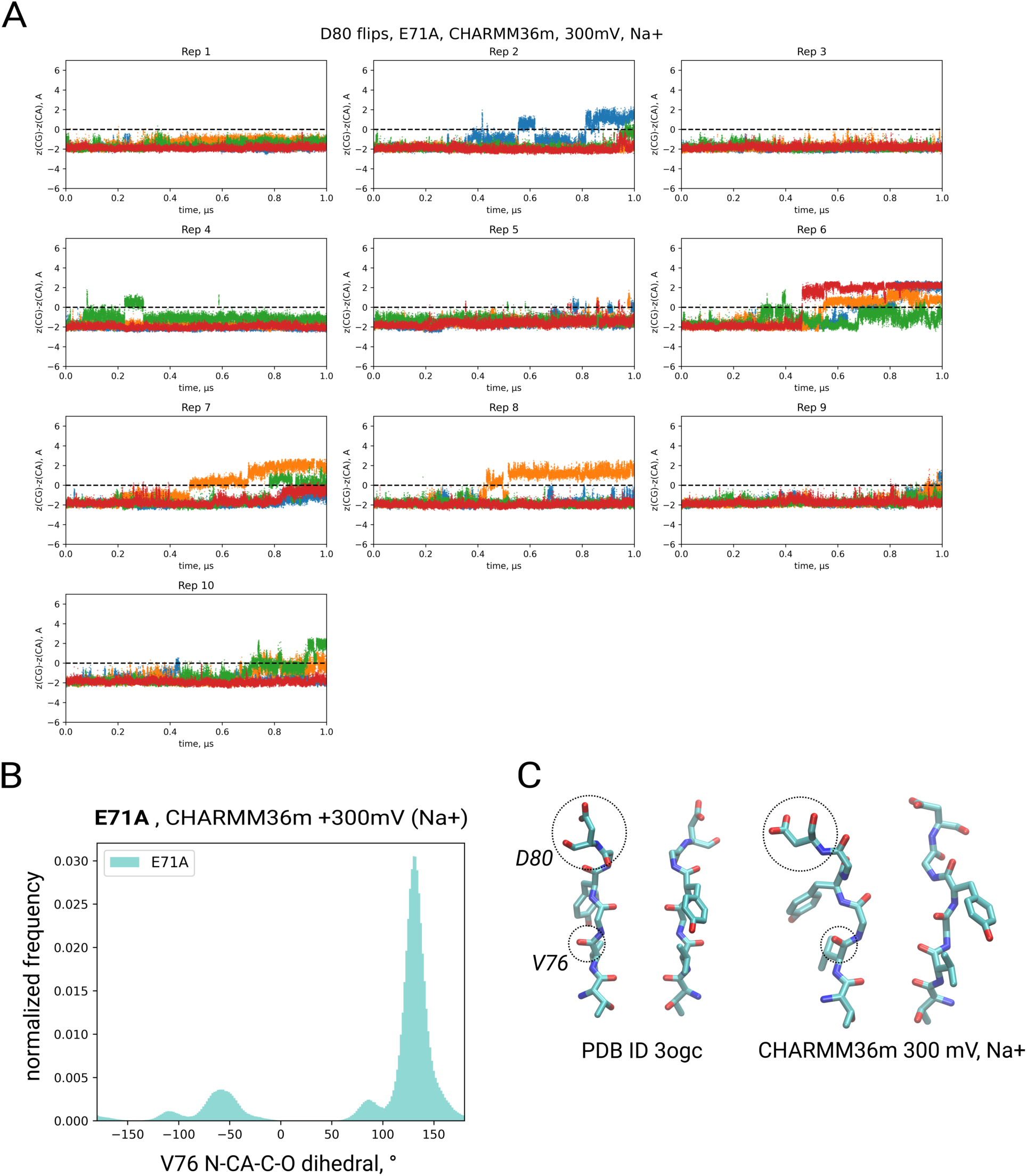
(*A*) Traces of differences between z positions of CG and CA atoms of D80 for KcsA E71A, CHARMM36m, 300 mV, in presence of Na^+^ but not K^+^. Positions above the dashed line indicate a flipped D80. Colors correspond to different subunits. (*B*) Distribution of V76 N-CA-C-O dihedrals, the region from −130 to −50 corresponds to a flipped state. (*C*) The SF of KcsA E71A in the crystal structure obtained in Na^+^-only conditions (left), and an example conformation with some D80 and V76 flipped as seen in our simulations (right).

**SI Fig. 6.**
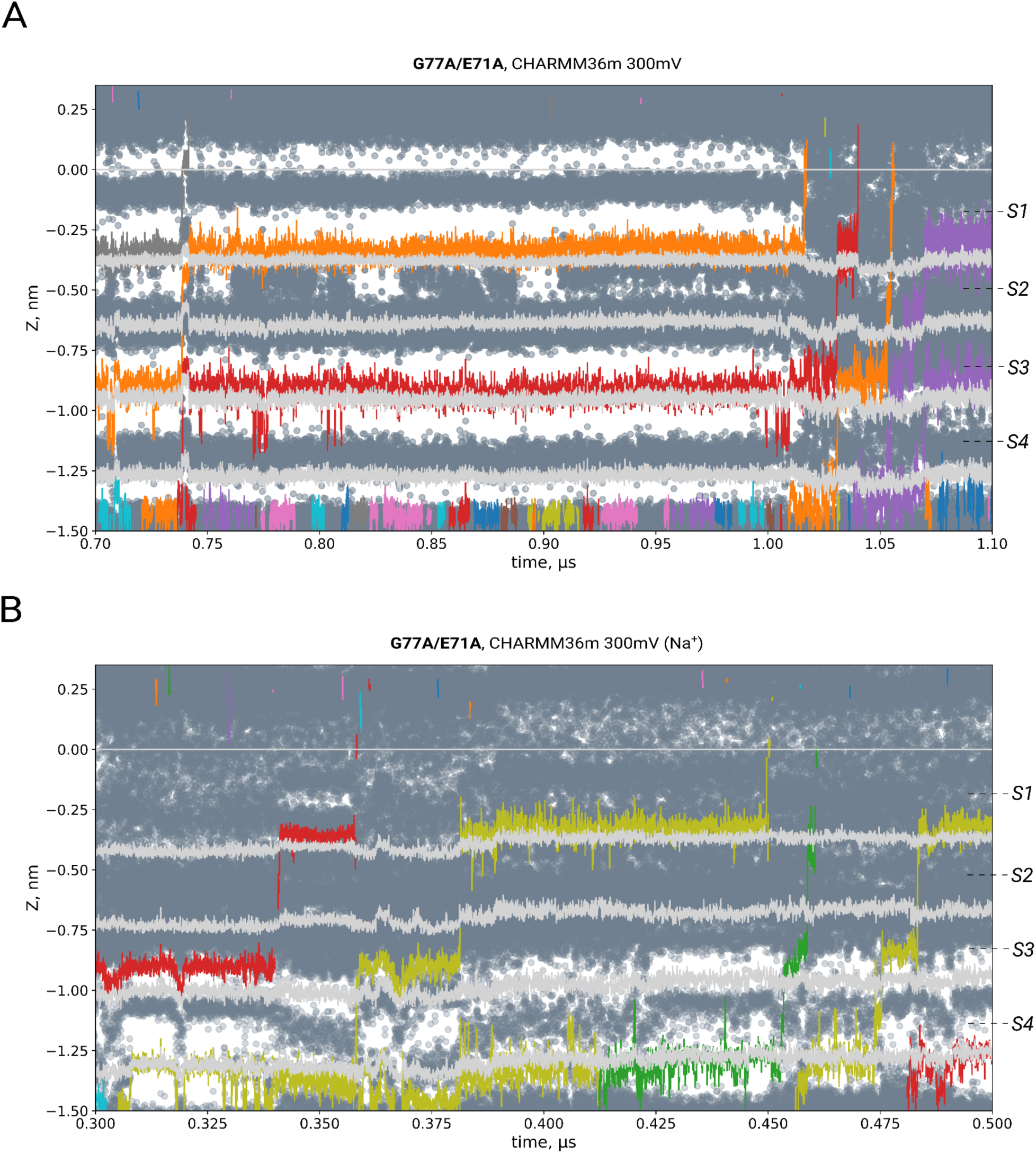
(*A*) K^+^ (colored lines), water (dark gray scatter) and backbone carbonyl oxygens’/ threonine hydroxyls traces (light gray lines) in the SF of G77A/E71A in CHARMM36m show permeation as jumps between KWKW-like configurations, without an intermediate WKWK. (*B*) Na^+^ ion permeation in G77A/E71A in CHARMM36m.

**SI Fig. 7.**
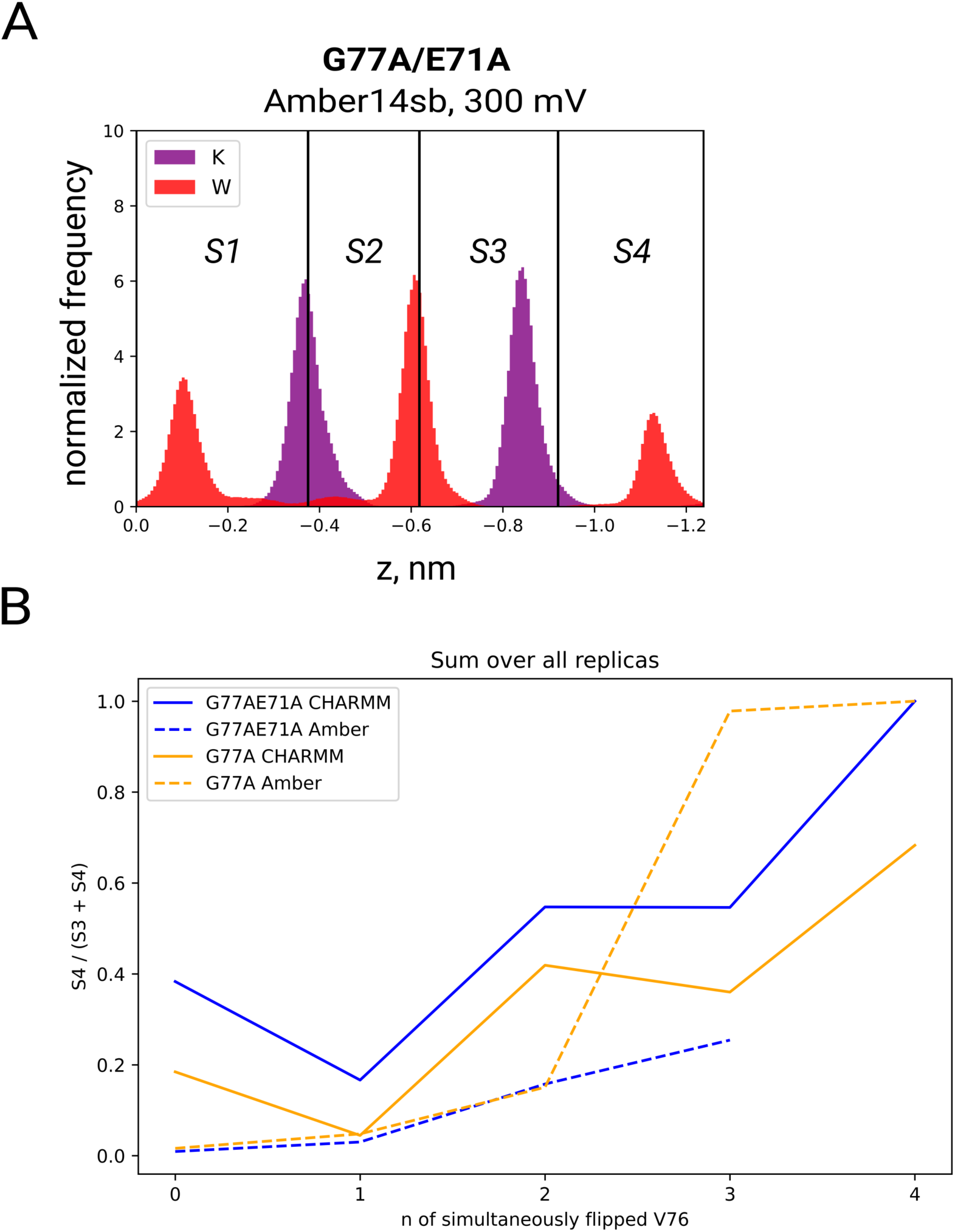
(*A*) Distribution of K^+^ and water in the SF of G77A/E71A, Amber14sb (all simulation replicas at 300 mV). (*B*) Fraction of K^+^ in S4 opposed to S3 for n (0-4) simultaneously flipped V76 carbonyls for G77A/E71A and G77A, showing the tendency of K^+^ to prefer the crystallographic S4 when more carbonyls are flipped.

**SI Fig. 8.**
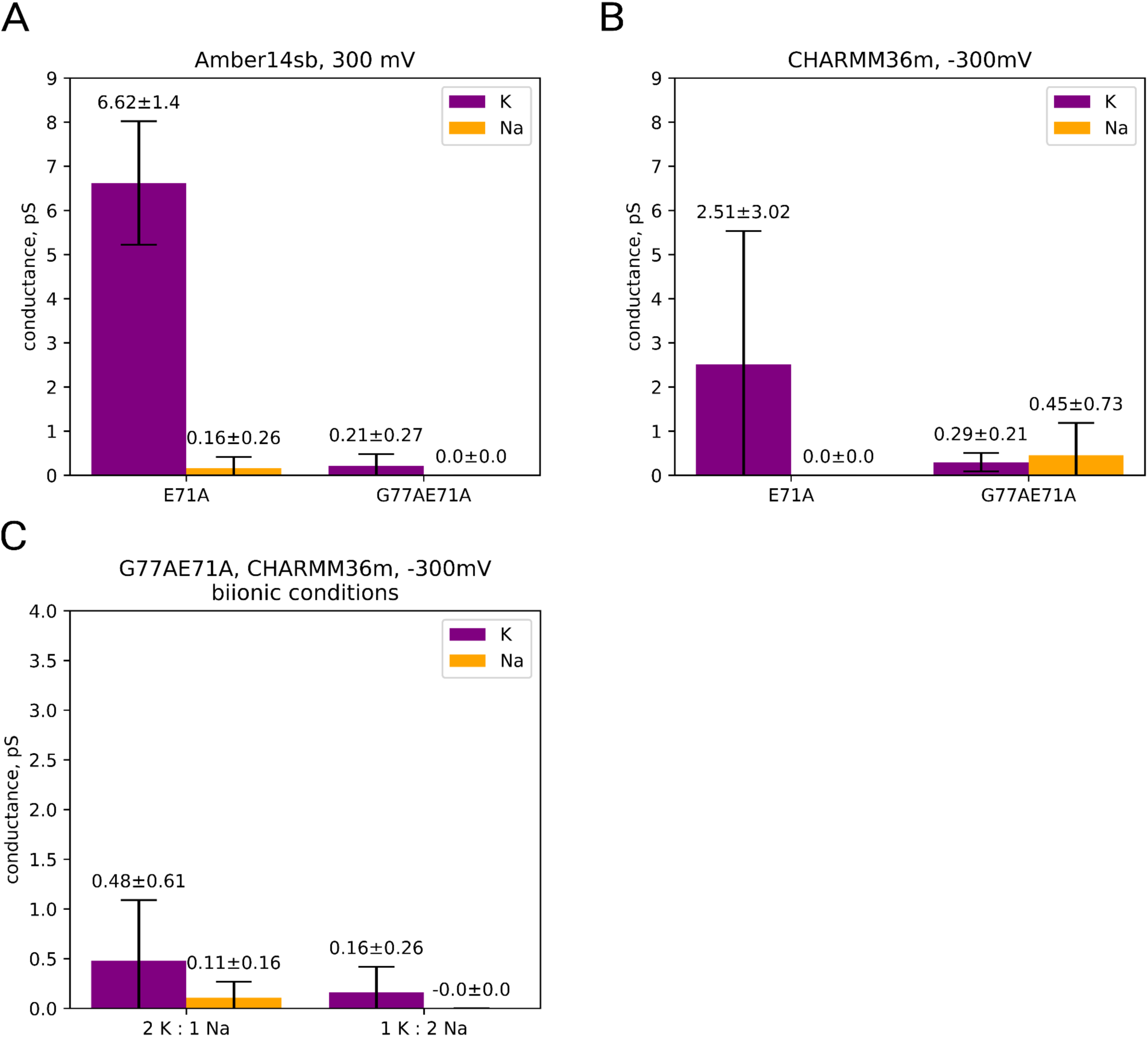
K^+^ and Na^+^ conductances of (*A*) E71A and G77A/E71A in Amber14sb, (*B*) E71A and G77A/E71A in CHARMM36m at negative voltage, (*C*) G77A/E71A in CHARMM36m at negative voltage when both K^+^ and Na^+^ are present at respective concentration ratios.

**SI Fig. 9.**
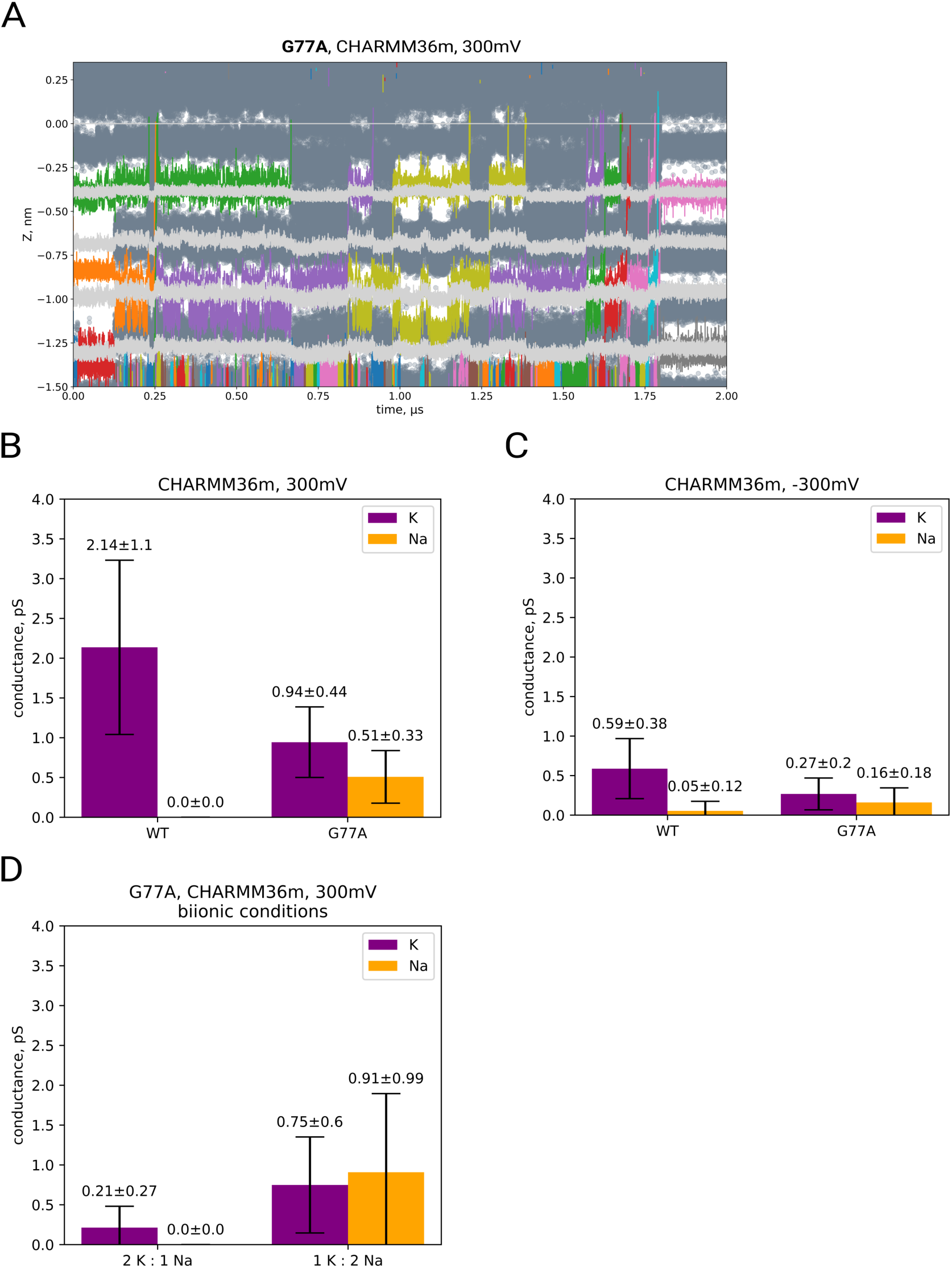
(*A*) Co-permeation of K^+^ (colored lines) and water (dark gray scatter) in the SF of G77A in CHARMM36m. Positions of backbone carbonyl oxygens and threonine hydroxyls are shown as light gray lines. K^+^ and Na^+^ conductances of WT and G77A in CHARMM36m at (*B*) positive and (*C*) negative voltages. (*D*) K^+^ and Na^+^ conductances of G77A in CHARMM36m at positive voltage and with both ions present at given concentration ratios.

**SI Fig. 10.**
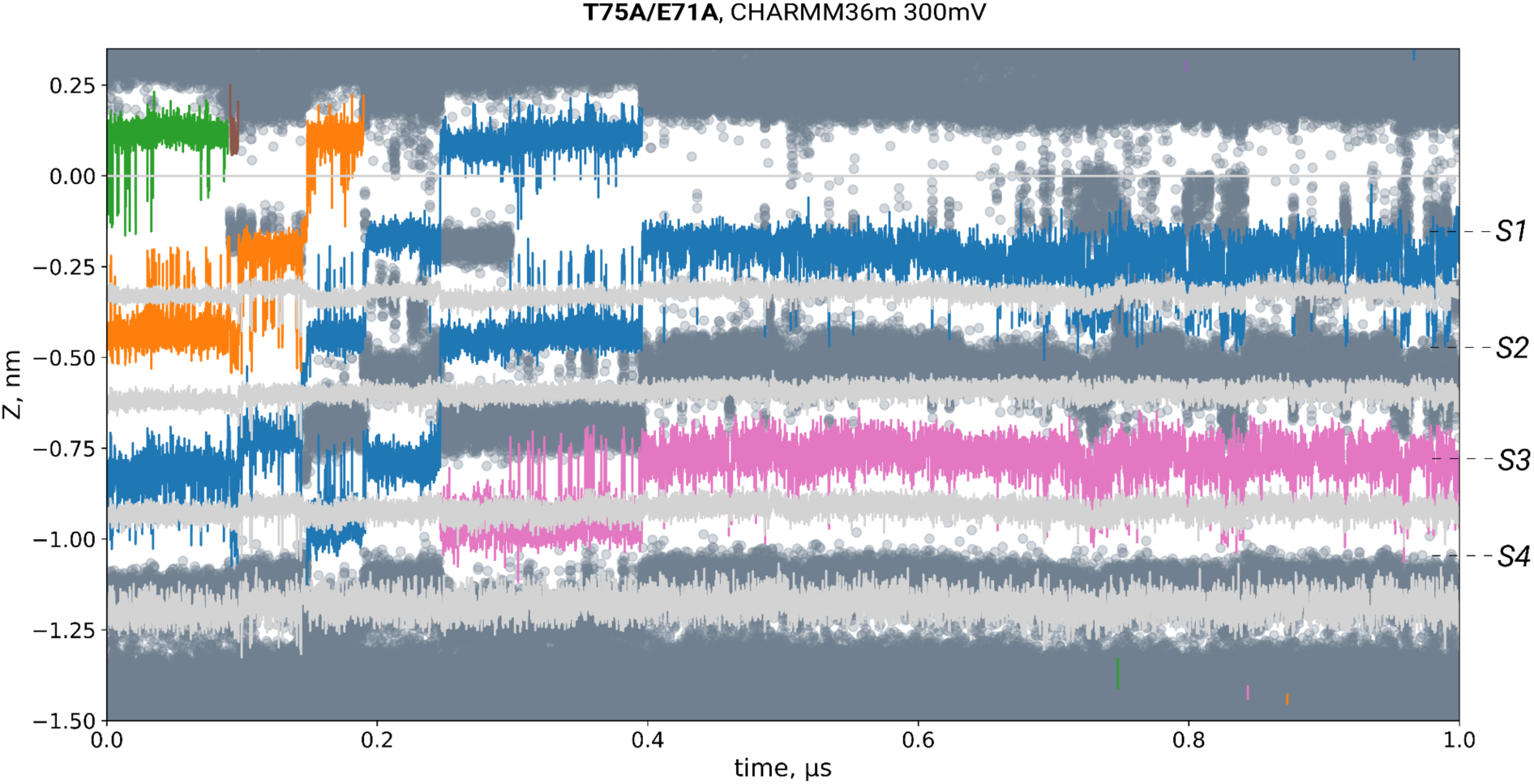
Co-permeation of K^+^ (colored lines) and water (dark gray scatter) in the SF of T75A/ E71A in CHARMM36m. Positions of backbone carbonyl oxygens and one of the hydrogens on the CB atom of A75 (instead of the T75 hydroxyls) are shown as light gray lines.

**SI Fig. 11.**
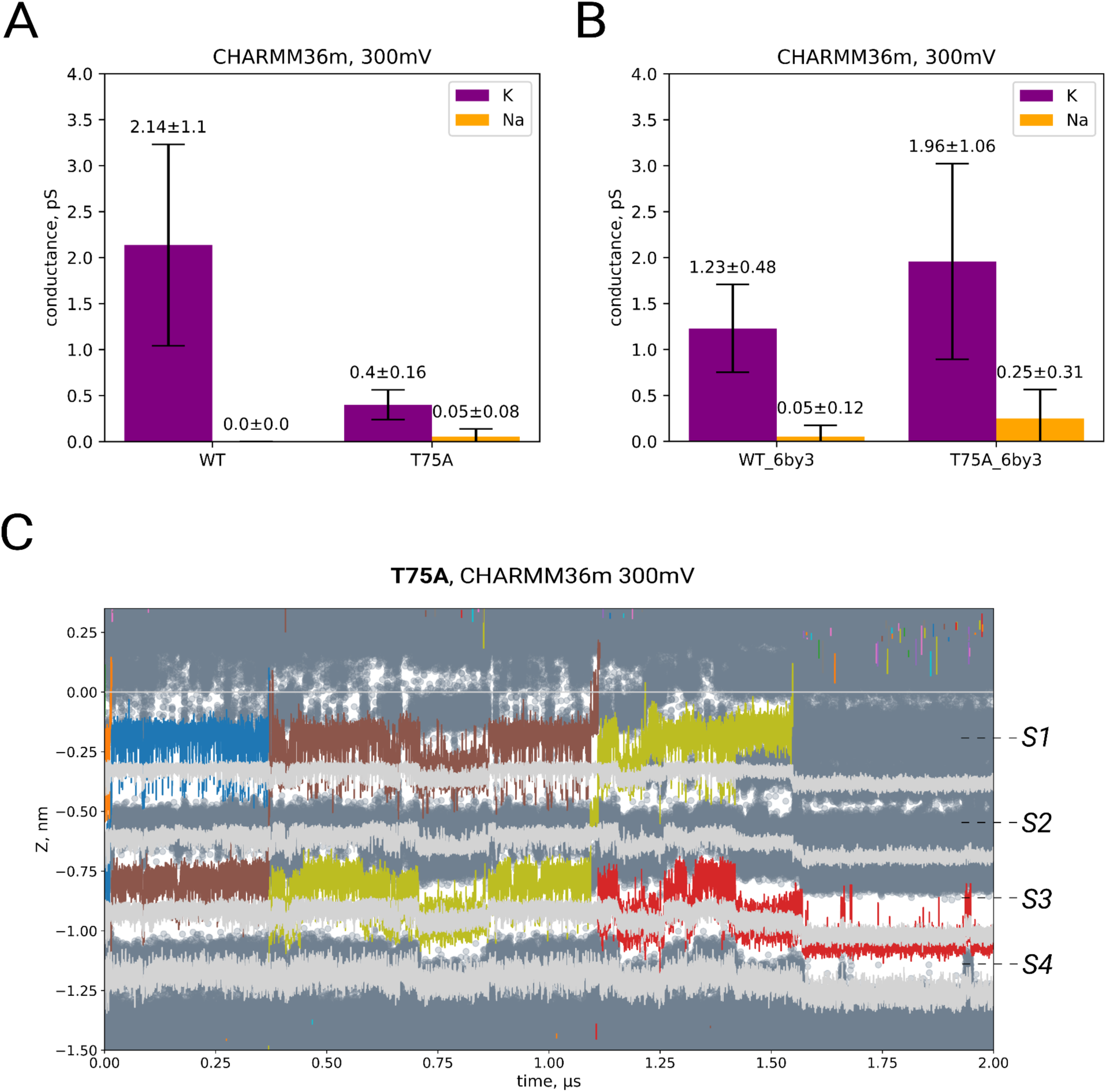
(*A*) K^+^ and Na^+^ conductances of WT and T75A at 300 mV in CHARMM36m from the main simulation set, and (*B*) derived from the crystal structure of the open state of KcsA T75A (PDB ID 6by3 (24)). (*C*) Traces of K^+^ (colored lines), water (dark gray scatter) and backbone carbonyl oxygens/one of the hydrogens on the CB atom of A75 instead of the T75 hydroxyls (light gray lines) in the SF of T75A from the main simulation set in CHARMM36m.

**SI Fig. 12.**
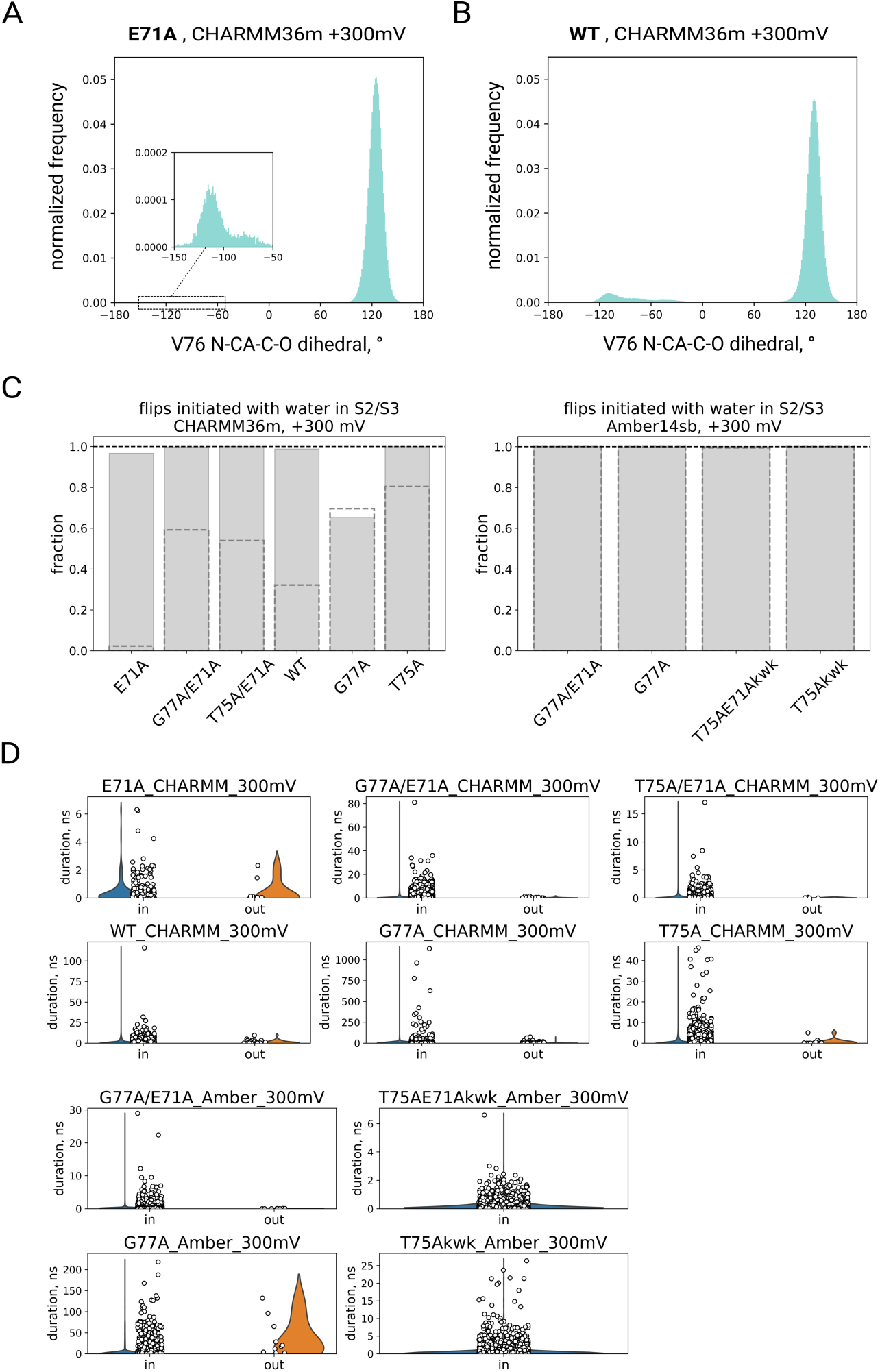
Both E71A (*A*) and WT KcsA (*B*) show low frequency of V76 carbonyl flipping, as shown by distributions of V76 N-CA-C-O dihedrals. (*C*) Solid bars represent fractions of flipping events initiated when water was already present in SF sites S2/S3 out of all flipping events. A flipping event is defined as a continuous stretch of simulation where at least one V76 is flipped. Systems where flips weren’t detected are not shown. Dashed bars represent the fraction of frames with water in S2/S3 out of all simulation frames; as water is not always present in S2/S3 in CHARMM36m (*left*), but the majority of flips still happen from the water-filled SF, this suggests that water causes V76 flips. In Amber14sb, establishing a causal relationship was more difficult, as the SF always contained water molecules. (*D*) Distribution of durations of flipping events that started when water was already in S2/S3 (‘in’) and when no water was present in these sites (‘out’).

**SI Fig. 13.**
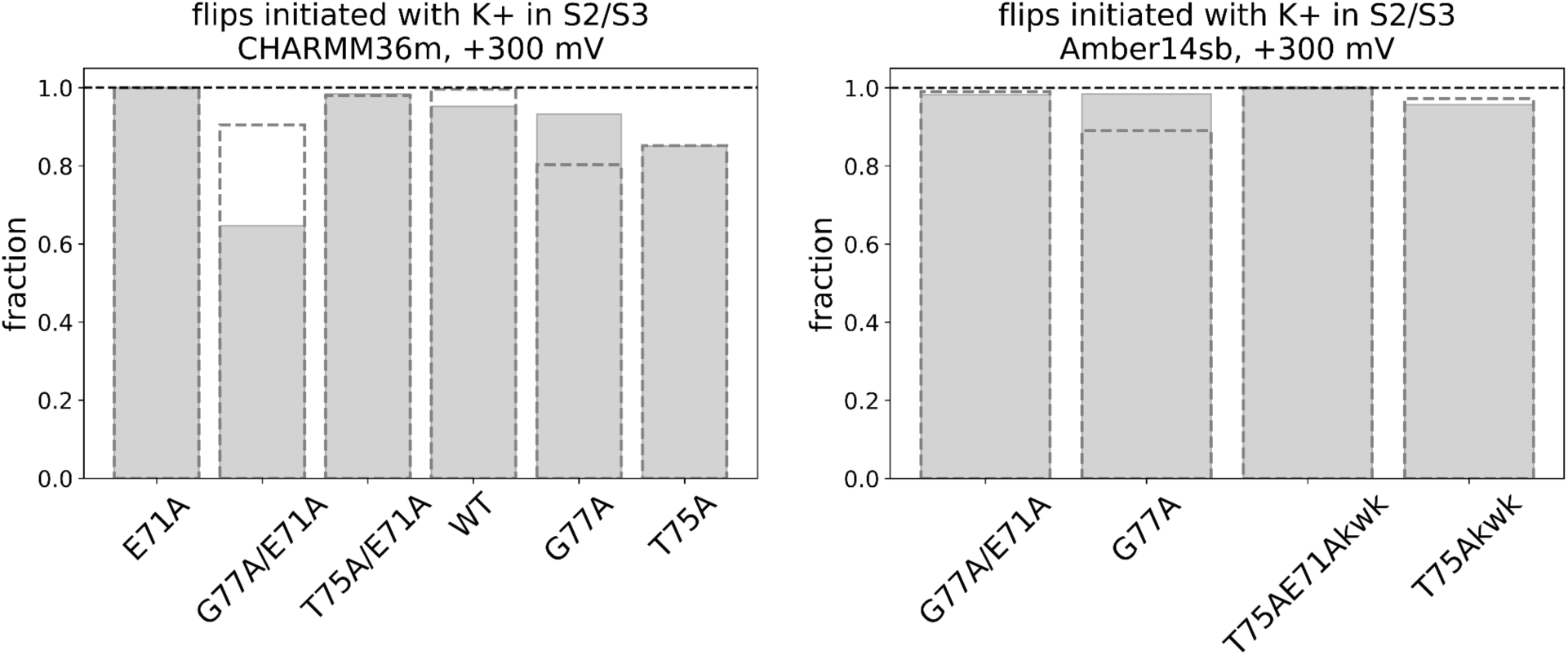
Fractions of flipping events initiated when K^+^ was present in SF sites S2/S3 out of all flipping events. Systems where flips haven’t been detected are not shown. In Amber14sb flips occurred only when soft knock-on-like starting SF configurations were used (‘kwk’). Bars with dashed edges represent the fraction of frames where K^+^ were in S2/S3, out of all simulation frames.

**SI Fig. 14.**
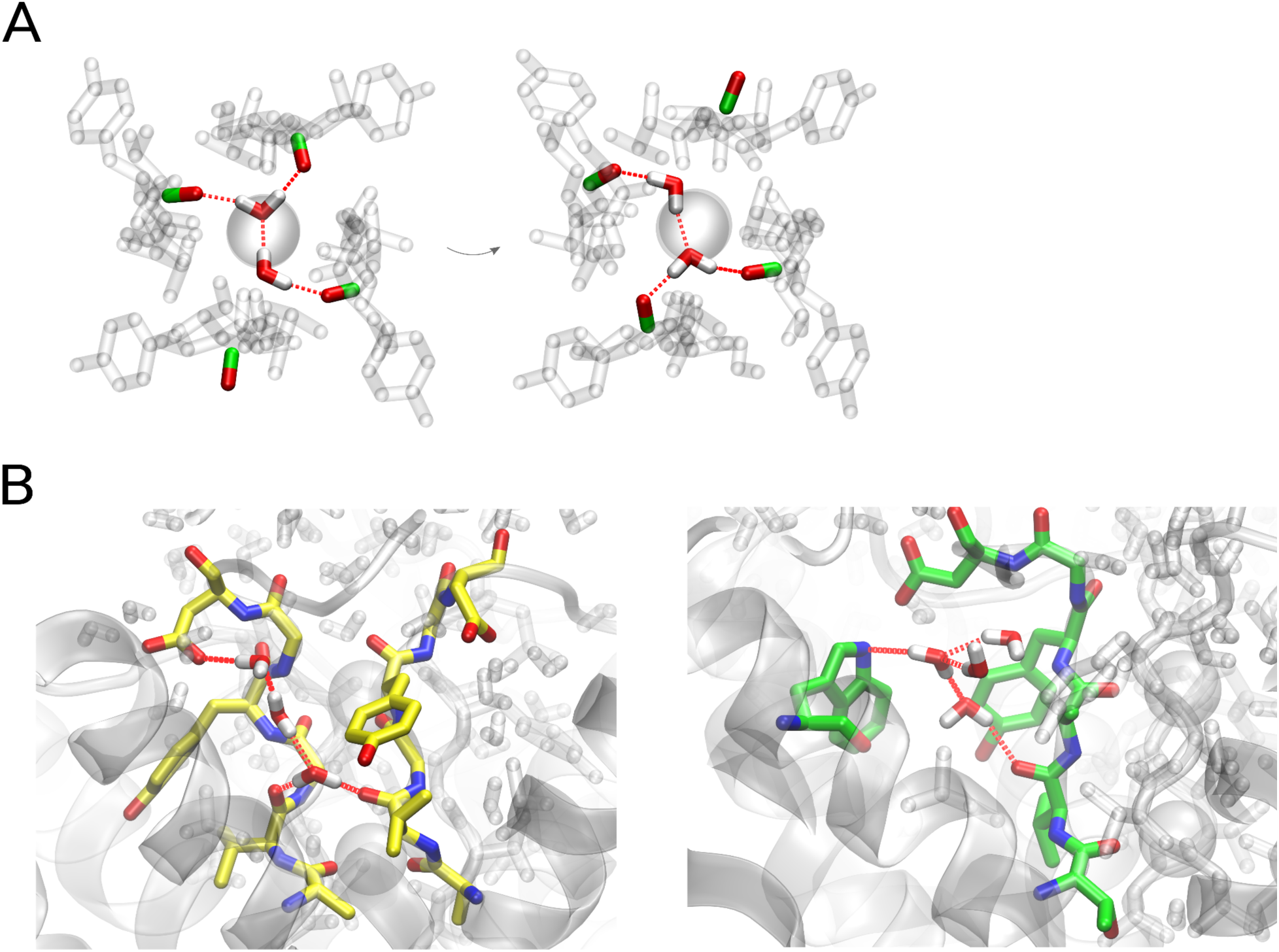
(*A*) Possible mechanism of stabilization of flipped and non-flipped V76 carbonyls through specific hydrogen bond networks when water molecules are present in S2 or S3. Water molecules and V76 carbonyls are opaque. Snapshots show a rearrangement of the network accompanied by a flipping of one carbonyl, and unflipping of another. (*B*) Stabilization of flips by hydrogen bond networks involving water molecules behind the SF in T75A/E71A (left) and G77A/E71A (right).

**SI Fig. 15.**
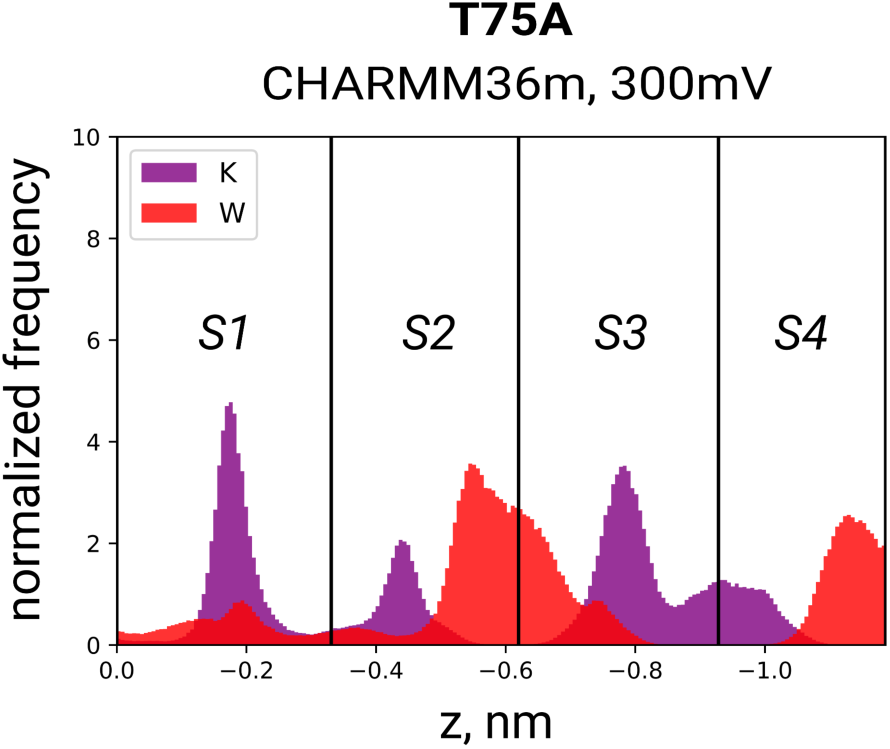
Distribution of K^+^ and water in the SF of T75A in CHARMM36m (all simulation replicas at 300 mV).

**SI Fig. 16.**
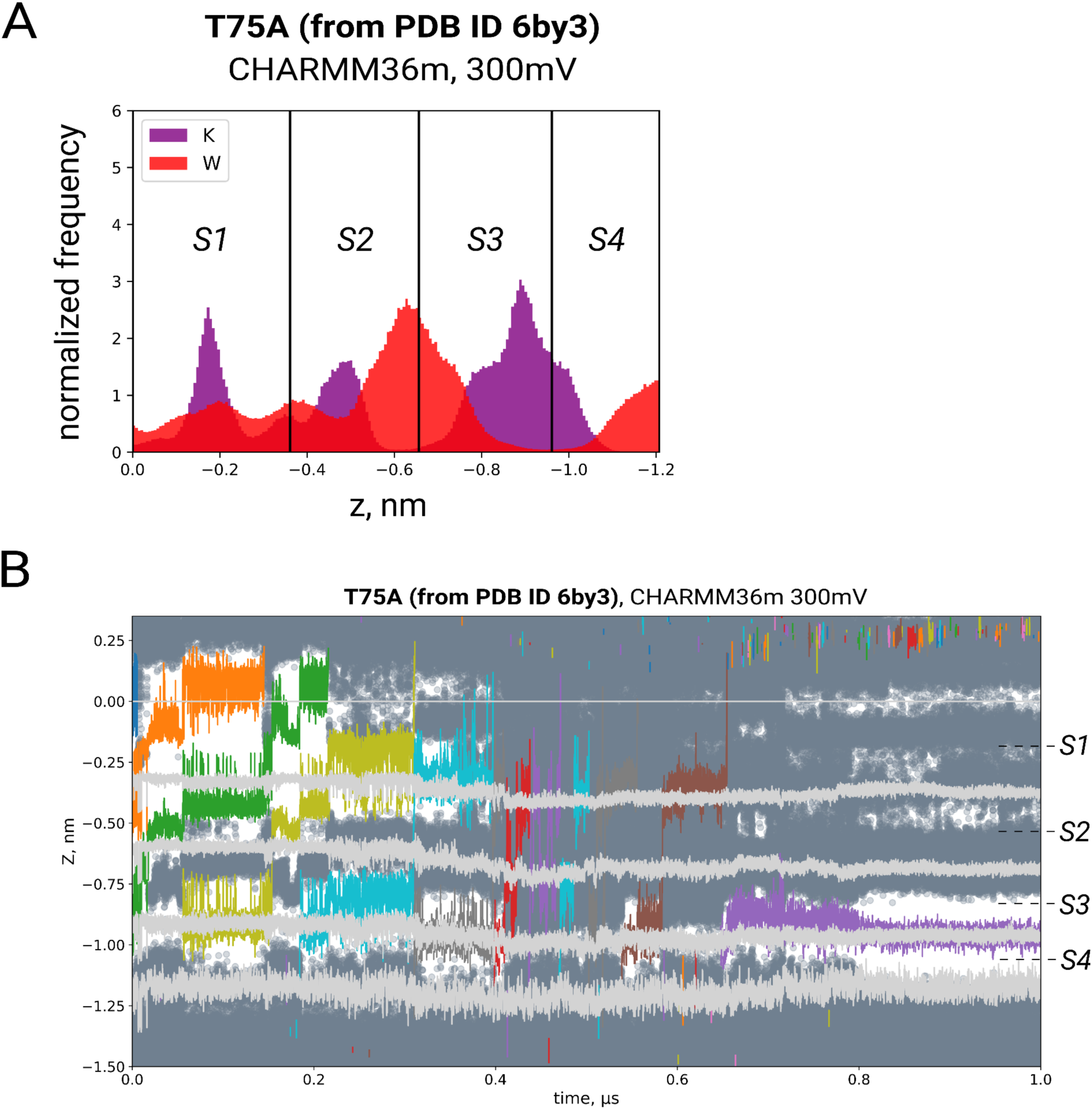
(*A*) K^+^/water distribution and (*B*) traces showing K^+^ (colored lines) and water (gray scatter) permeation in the SF (backbone carbonyls and one of the hydrogens on the CB atom of A75 (instead of the T75 hydroxyls) are represented by light gray lines) of T75A built from the crystal structure of the open state of KcsA T75A (PDB ID 6by3 (24)).

**SI Fig. 17.**
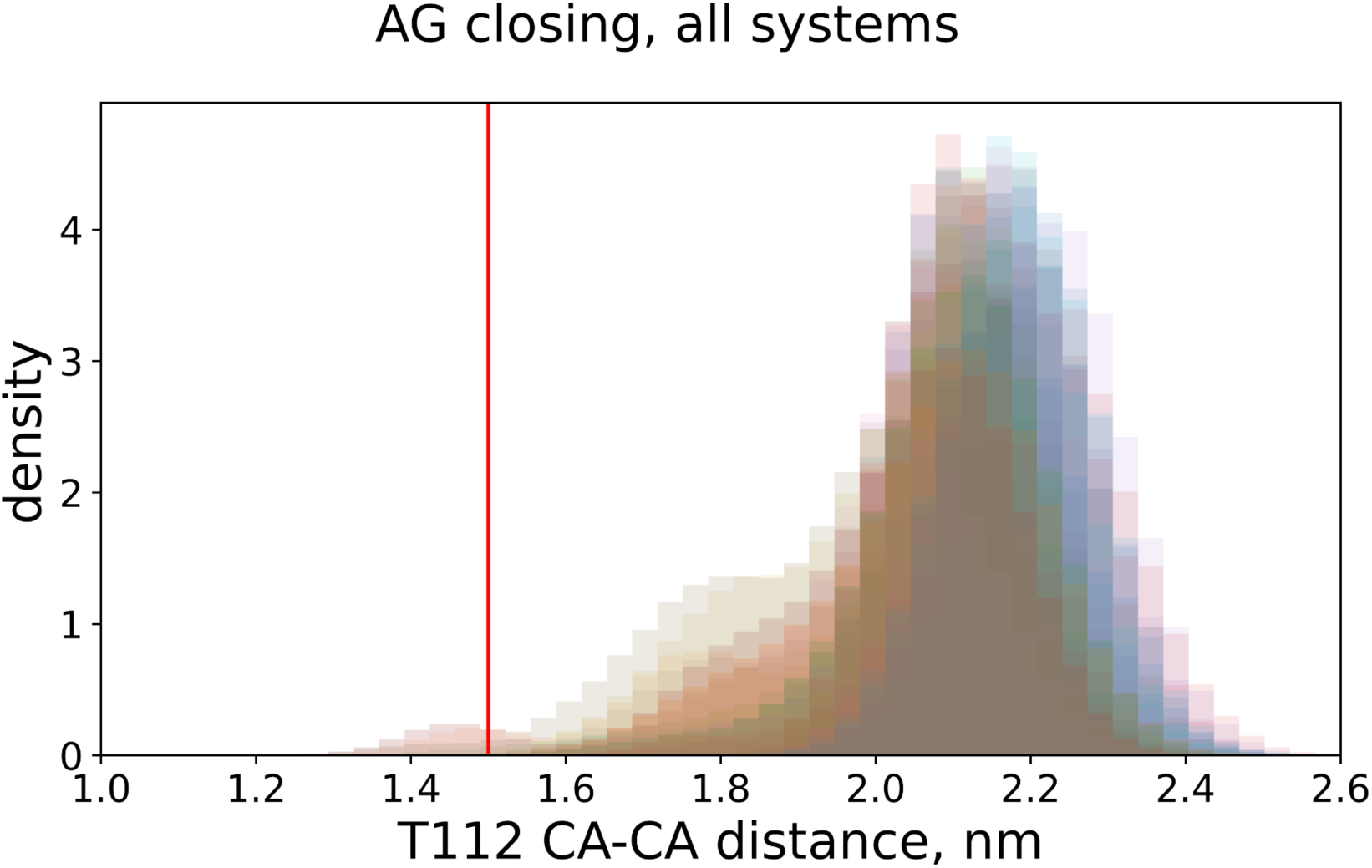
Distribution of distances between T112 CA atoms of opposite subunits in all systems and conditions as a measure of the state of the activation gate. Red line indicates the activation gate opening in the partially open state (∼1.5 nm, PDB ID 3fb5 (70)) below which we considered the gate to be closed. The channel remained in the open state for the majority of simulation time.

---

**SI Table 1.**
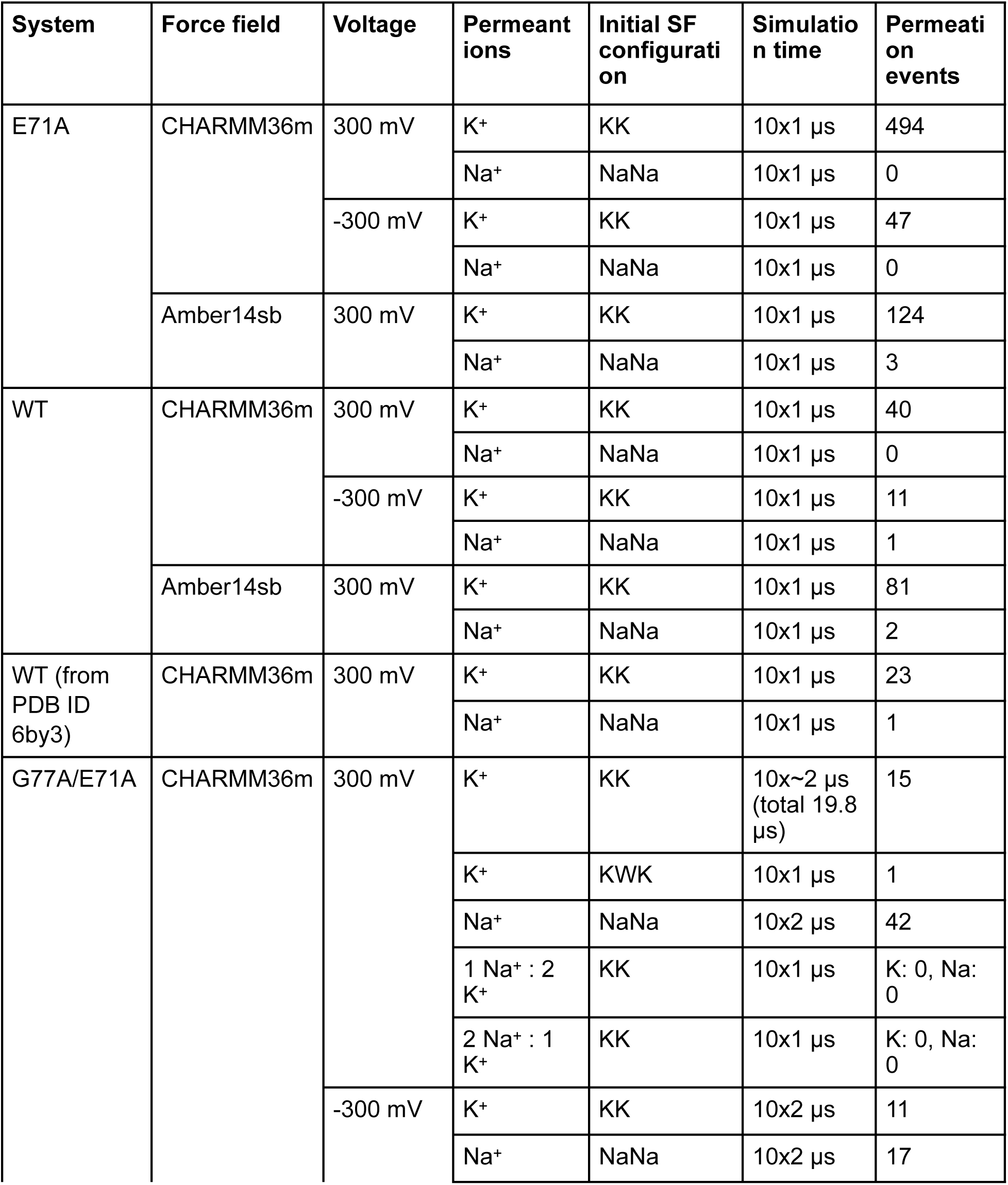
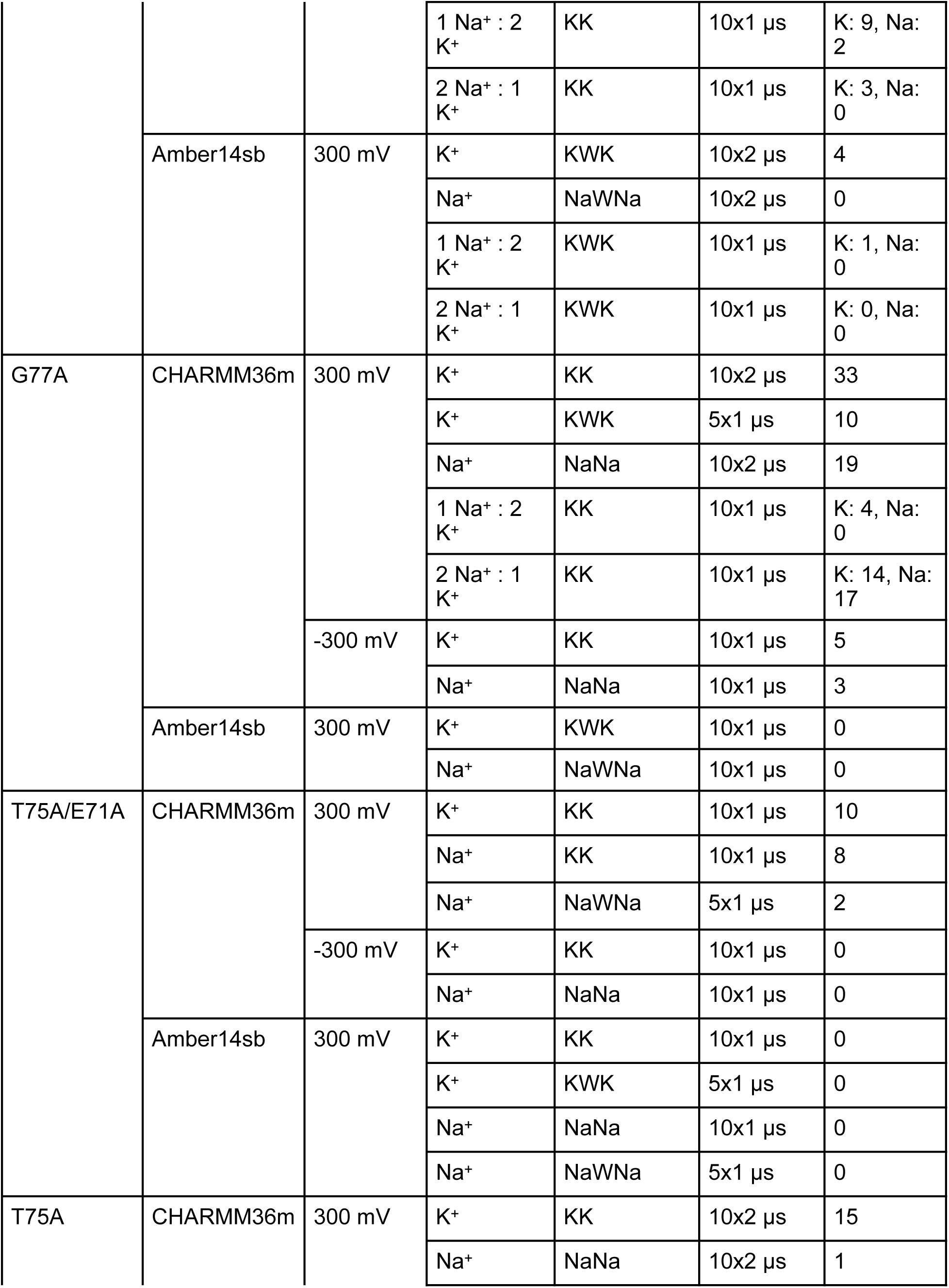
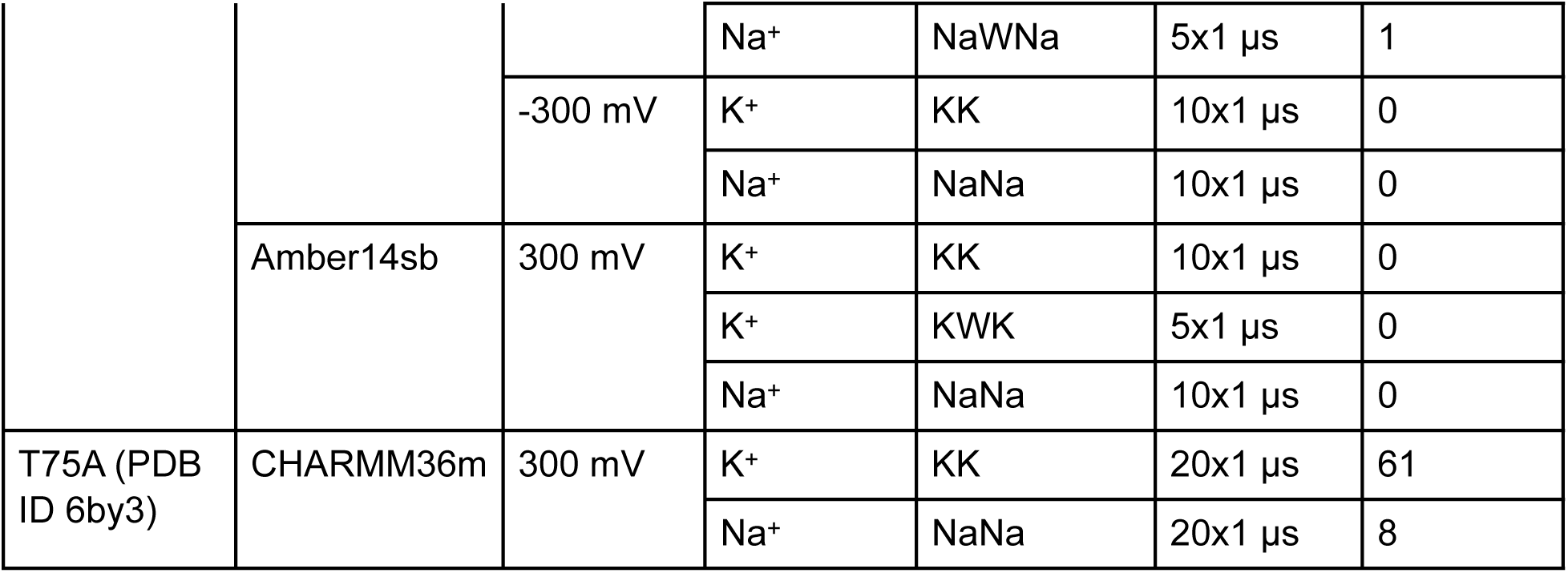
List of all simulated systems. ‘KK’ and ‘KWK’ (or ‘NaNa’ and ‘NaWNa’) indicate direct or soft knock-on-like initial configurations containing K^+^ or Na^+^ ions in the SF, respectively. For simulations in biionic conditions, ratios of Na^+^ and K^+^ concentrations are given. Unless stated, all systems are built from PDB ID 5vk6.

